# Vesiculation pathways in clathrin-mediated endocytosis

**DOI:** 10.1101/2024.08.13.607731

**Authors:** Xinran Wang, Julien Berro, Rui Ma

**Author notes:** **For correspondence:** (FMS); (FS).

## Abstract

During clathrin-mediated endocytosis, a patch of flat plasma membrane is internalized to form a vesicle. In mammalian cells, how the clathrin coat deforms the membrane into a vesicle remains unclear and two main hypotheses have been debated. The “constant area” hypothesis assumes that clathrin molecules initially form a flat lattice on the membrane and deform the membrane by changing its intrinsic curvature while keeping the coating area constant. The alternative “constant curvature” hypothesis assumes that the intrinsic curvature of the clathrin lattice remains constant during the formation of a vesicle while the surface area it covers increases. Previous experimental studies were unable to unambiguously determine which hypothesis is correct. In this paper, we show that these two hypotheses are only two extreme cases of a continuum spectrum if we account for the free energies associated with clathrin assembly and curvature generation. By tracing the negative gradient of the free energy, we define vesiculation pathways in the phase space of the coating area and the intrinsic curvature of clathrin coat. Our results show that, overall, the differences in measurable membrane morphology between the different models are not as big as expected, and the main differences are most salient at the early stage of endocytosis. Furthermore, the best fitting pathway to experimental data is not compatible with the constant-curvature model and resembles a constant-area-like pathway where the coating area initially expands with minor changes in the intrinsic curvature, later followed by a dramatic increase in the intrinsic curvature and minor change in the coating area. Our results also suggest that experimental measurement of the tip radius and the projected area of the clathrin coat will be the key to distinguish between models.

## Introduction

Clathrin-mediated endocytosis (CME) is a fundamental cellular process to transport lipids, membrane proteins and extracellular cargo molecules into the cell (***McMahon and Boucrot, 2011***; ***Sorkin and Puthenveedu, 2013***; ***Lu et al., 2016***; ***Kaksonen and Roux, 2018***; ***Lacy et al., 2018***; ***Mettlen et al., 2018***). In mammalian cells, a small patch of flat plasma membrane is shaped into a spherical vesicle when CME occurs (***Avinoam et al., 2015***). Clathrin molecules are essential for the membrane remodelling process. They are made of three subunits that form a triskelion, which further assemble into a cage-like structure *in vitro* (***Musacchio et al., 1999***; ***Shraiman, 1997***). The minimum cages contain 16 polygons (***Fotin et al., 2004***) and the most commonly observed ones are semi-regular icosahedral cages (***Cheng et al., 2007***; ***Dannhauser and Ungewickell, 2012***; ***Heuser, 1980***). Two main hypotheses are under debate regarding how the clathrin coat scaffolds the flat membrane into a spherical vesicle *in vivo* (***Avinoam et al., 2015***; ***Chen and Schmid, 2020***; ***Frey and Schwarz, 2020***; ***Kaksonen and Roux, 2018***; ***Scott et al., 2018***). The constant area model asserts that the clathrin molecules initially polymerize into a flat lattice with a regularly arranged hexagonal structure, and later reorganization of the bonds between adjacent clathrins results in the formation of pentagons in the hexagonal lattice, which in turn leads to curvature generation of the clahtrin coat (***Fotin et al., 2004***; ***Heuser and Anderson, 1989***) (***Figure 1***a). Adhesion of clathrin molecules with the substrate has been suggested to contribute a flattening force that prevents curvature generation. Release of the flattening force therefore could induce curvature of the clathrin coat with preloaded pentagons (***Sochacki et al., 2021***). The alternative constant curvature model asserts that the intrinsic curvature of the elements of the lattice is kept constant during the assembly and expansion of the lattice, therefore curvature generation occurs from the very beginning of clathrin assembly (***Figure 1***b).

**Figure 1.**
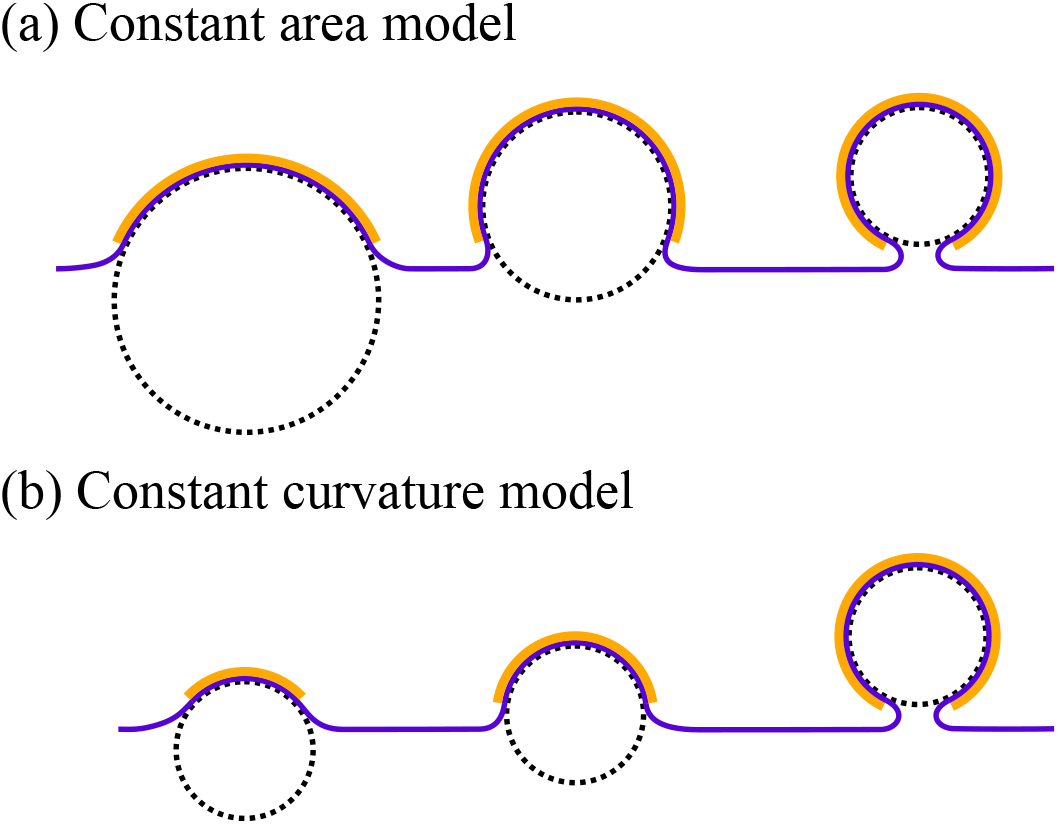
Schematic illustrations of the constant area model (a) and the constant curvature model (b) for CME. Blue: plasma membrane, yellow: clathrin coat, black dashed line: curvature of the clathrin coat

In order to distinguish between the two models, the dynamics of clathrin assembly and the geometry of membrane shapes are needed. Fluorescence microscopy, including light sheet and MINFLUX, has revealed the assembly dynamics of the clathrin coat (***Aguet et al., 2016***; ***Ferguson et al., 2016***), as well as other proteins that participate in endocytosis (***Sirotkin et al., 2010***; ***Taylor et al., 2011***; ***Kukulski et al., 2016***; ***Kaksonen et al., 2006***; ***Balzarotti et al., 2017***; ***Gwosch et al., 2020***; ***Schmidt et al., 2021***), while electron tomography has been able to resolve membrane shapes during endocytosis (***Avinoam et al., 2015***; ***Kukulski et al., 2012***). However, neither of the methods can capture both spatial and temporal information at the same time. Under conventional fluorescence microscopy, the clathrin-coated pits appear as diffraction-limited spots due to their small size, which is typically ~ 30 − 150nm in mammals (***McMahon and Boucrot, 2011***) and yeast (***Kukulski et al., 2012***), and shape information of the membrane is completely lost (***Cocucci et al., 2012***; ***Loerke et al., 2009***; ***Picco et al., 2015***; ***Sirotkin et al., 2010***; ***Kural and Kirchhausen, 2012***). On the other hand, super-resolution fluorescence microscopy has been able to reveal the protein organization at the endocytic pit (***Mund et al., 2018***; ***Cocucci et al., 2012***; ***Arasada et al., 2018***) and to reconstruct the shape of the clathrin coat (***Sochacki et al., 2017***; ***Scott et al., 2018***; ***Mund et al., 2023***) from averaging over ensembles of endocytic sites. Correlative light and electron microscopy (CLEM) method has exploited the fluorescence of fiducial markers to locate endocytic sites while resolving membrane shapes using electron tomography. However, both super-resolution and CLEM requires sample fixation, therefore, one can identify multiple endocytic sites at the same time and perform the average, yet unable to trace a single endocytic site over time. The temporal information is nevertheless lost.

As a result of the incomplete information obtained by existing experimental methods, both hypotheses have experimental support. Experiments that apply electron microscopy to resolve the membrane shapes of endocytic pits favor the constant area model (***Avinoam et al., 2015***; ***Bucher et al., 2018***; ***Sochacki and Taraska, 2019***; ***Sochacki et al., 2021***). However, super-resolution imaging combined with analysis of the fluorescence intensity of the clathrin coat is inclined towards the constant curvature model (***Willy et al., 2021***). In addition, it was argued that the energetic cost of bond reorganization in a regular hexagonal lattice in the constant area model may be too large to be fulfilled (***Frey et al., 2020***; ***Frey and Schwarz, 2020***; ***Kirchhausen et al., 2014***).

Extensive theoretical efforts have been dedicated to model membrane morphology during endocytosis (***Fu and Johnson, 2023***), of which molecular dynamics simulations (***Varga et al., 2020***) and continuum mechanics (***Hassinger et al., 2017***; ***Ma and Berro, 2021***; ***Walani et al., 2015***; ***Agrawal and Steigmann, 2008***; ***Rangamani et al., 2013***) are two common approaches. Hybrid models were also broadly applied to gain higher resolution than continuum mechanics and lower computing expense than molecular dynamics (***Fu et al., 2019, 2021***). However, most theoretical investigations have focused on how mechanical properties, such as membrane tension and bending rigidity of the clathrin coat, influence the membrane morphology. The process of curvature generation is either neglected or taken for granted. Only few of them have addressed the difference between the constant area model and the constant curvature model ***Mund et al. (2023***).

In fact, the constant curvature and constant area models are only two extreme models for clathrin assembly during endocytosis and any change in area or curvature are possible at any time point during endocytosis. In this paper, we extend the classic Helfrich theory for membrane deformation to incorporate energy terms associated with clathrin assembly and curvature generation, and compare geometric features calculated by theory with those extracted from experimental data. The negative gradient of the total free energy defines a pathway that neither fits the constant area model nor the constant curvature model. We find that a pathway that is close to the constant area model fits electron tomograms of the endocytic pits the best. Our study also offers experimental suggestions to distinguish between the two main hypotheses.

### Models and methods

We model the membrane patch of the CCP (clathrin-coated pit) as a surface which is rotationally symmetric with respect to the *z*-axis. The shape of the membrane is parameterized with the meridional curve {*r*(*s*), *z*(*s*)}, where *s* denotes the arc length along the curve. The bending energy of the membrane (together with the clathrin coat) assumes the Helfrich model (***Helfrich, 1973***)

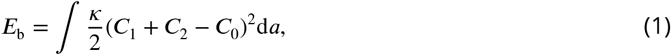

where *κ* denotes the bending rigidity of the CCP, *C*_1_ and *C*_2_ denote the principal curvatures of the surface, *C*_0_ denotes the intrinsic curvature of the membrane induced by the clathrin coat. To model a finite area of the clathrin coat, we assume the intrinsic curvature *C*_0_ spatially varies as

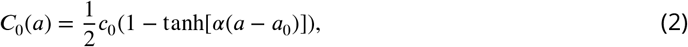

where *a* denotes positions on the membrane. Here, we choose *a* to be the surface area calculated from the tip of the membrane, and *C*_0_ equals *c*_0_ for area *a* < *a*_0_, and rapidly drops to zero when *a* > *a*_0_. The parameter α controls the sharpness of the drop. In the constant area model, we vary the intrinsic curvature *c*_0_ but keep the coating area *a*_0_ constant, while in the constant curvature model, we vary the coating area *a*_0_ but keep the intrinsic curvature *c*_0_ constant. As a result of the clathrin coat, the bending rigidity *κ* also varies as

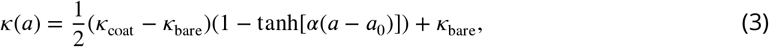

where *κ*_coat_ and *κ*_bare_ denote the bending rigidity of the clathrin-coated membrane and bare membrane, respectively. Besides the bending energy, the membrane tension contributes to the free energy in the form of

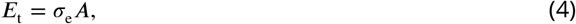

where *σ*_e_ denotes the membrane tension at the base and *A* denotes the surface area of the membrane patch within a fixed radius of *R*_b_. The membrane tension *σ*_e_ and bending rigidity *κ* define a characteristic length 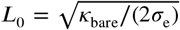 (***Derényi et al., 2002***). The total free energy *E*_tot_ = *E*_b_ + *E*_t_ is a functional of the membrane shape. We emphasize that the *constant area model* refers to the assumption that the clathrin-coated area *a*_0_ remains fixed. Meanwhile, the membrane tension *σ*_*e*_ at the base is held constant, allowing the total membrane area *A* to vary in response to deformations induced by the clathrin coat. Given a coating area *a*_0_ and an intrinsic curvature *c*_0_, we numerically solve the variational equations of the energy functional to obtain membrane shapes that minimizes *E*_tot_. This approach is based on the assumption that the membrane remains in mechanical equilibrium throughout the process of clathrin assembly, which is justified by the disparity between the timescales of membrane relaxation and clathrin assembly. The characteristic relaxation time of a lipid membrane is given by 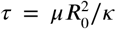, where *µ* ≈ 5 × 10^−9^ N ⋅ s ⋅ m^−1^ is the membrane viscosity ***Arroyo and DeSimone (2009)***, *R*_0_ ≈ 50 nm is the vesicle size ***Avinoam et al. (2015***), and *κ* ≈ 20 *k*_*B*_*T* is the bending rigidity ***Yuan et al. (2021***). Substituting these values yields a relaxation time of *τ* ≈ 1.5 × 10^−4^ s, which is several orders of magnitude shorter than the clathrin assembly timescale which is approximately one minute. Therefore, it is reasonable to assume that the membrane rapidly equilibrates and remains in mechanical equilibrium during the assembly process. More detailed descriptions of the model can be found in ***Appendix 1***.

We rescale lengths by the characteristic length *L*_0_ and energies by 2*πκ*_bare_. Dimensionless quantities are denoted with a bar. The dimensionless tension energy, defined as *Ē*_*t*_ = *E*_*t*_/(2*πκ*), simplifies to *Ē*_*t*_ = *Ā*/2, where *Ā* = *A*/(2*πL*^2^) is the dimensionless membrane area. Therefore, the dimensionless membrane tension becomes a constant 1/2. When presenting data, dimensionless quantities are shown on the left axes, and the corresponding dimensional values are shown on the right axes.

## Results

### Difference between the constant area model and the constant curvature model in terms of membrane morphology

Vesiculation requires assembly of a clathrin coat on the membrane, as well as curvature generation from the clathrin coat. A vesiculation process defines a pathway in the phase space (*a*_0_, *c*_0_) of the clathrin coat area *a*_0_ and the intrinsic curvature *c*_0_ of the coat. The constant area model and the constant curvature model are pathways that are made of a vertical line and a horizontal line. Besides these two extreme cases, there is a continuum spectrum of pathways with simultaneously increasing coating area *a*_0_ and intrinsic curvature *c*_0_ that could lead to vesiculation. Along the pathway, the membrane evolves from a flat shape to a dimple shape, and finally to an Ω-shape, as shown in ***Figure 2***. Hereafter we use the maximal tangential angle *ψ*_max_ of the membrane as an indicator of the progression of vesiculation - when the membrane is flat, *ψ*_max_ = 0^°^, and when the membrane becomes spherical, *ψ*_max_ = 180^°^. In our simulation, the neck becomes extremely narrow before *ψ*_max_ = 180^°^. Therefore, we consider vesiculation occurs when *ψ*_max_ reaches 150^°^ (***Figure 2***a).

**Figure 2.**
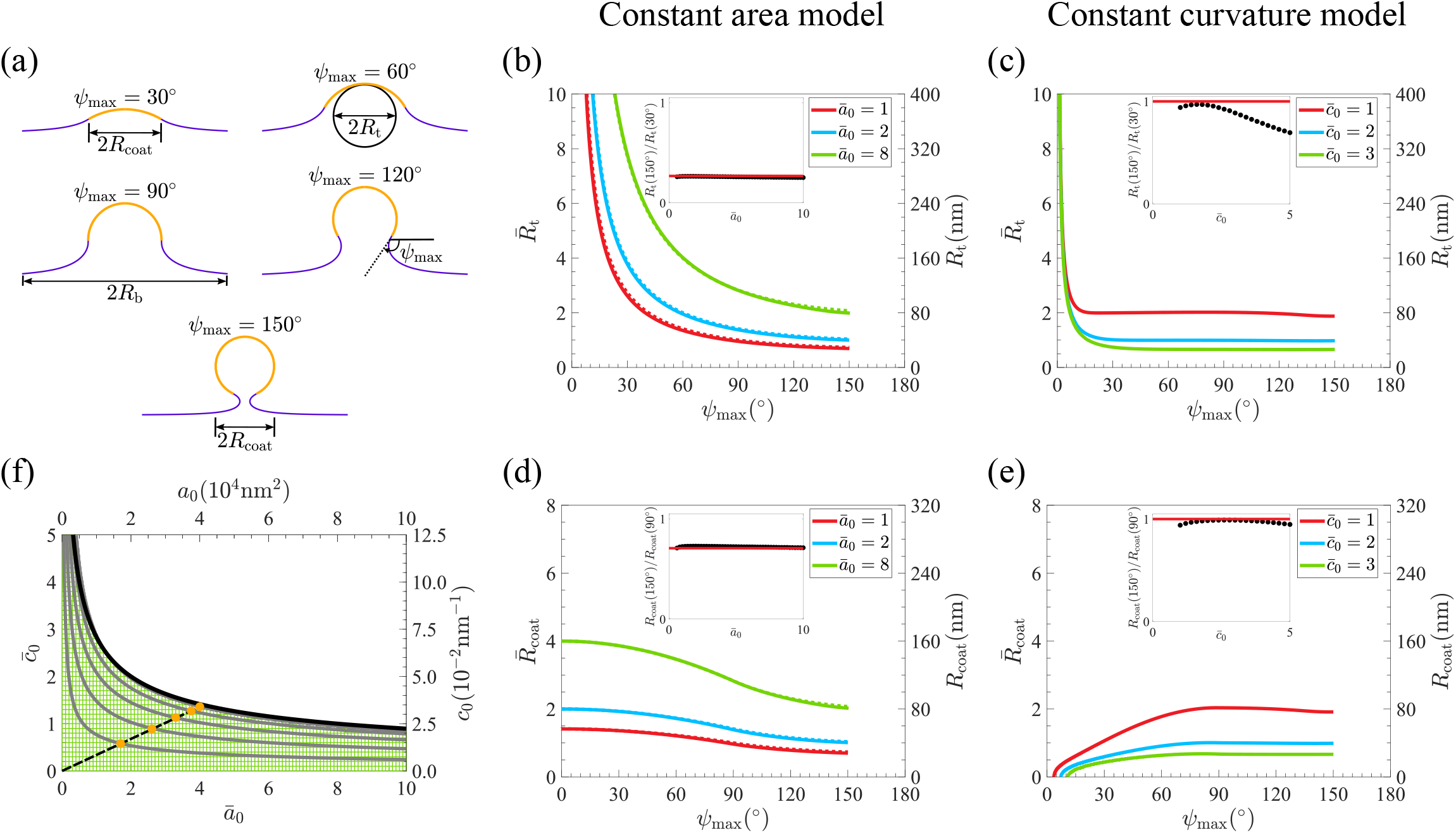
Evolution of membrane morphology and phase diagram of vesiculation in the coating area vs. intrinsic curvature (*a*_0_, *c*_0_) parameter space. (a) Membrane shapes at different stages of invagination and definition of some variables used in this paper. We define the distance from the axisymmetric axis to the edge of the coating area as *R*_coat_, the radius of the tangential curvature circle at the tip of the shape as *R*_t_, and the distance from axisymmetric axis to the boundary as *R*_b_. The maximum tangential angle of the cross section contour is *ψ*_max_. (b,c) Tip radius *R*_t_ vs. maximal angle *ψ*_max_ for the constant area model in (b) and for the constant curvature model in (c). Dotted lines in (b) denote the analytical solutions. Insets show the ratio of the tip radii *R*_t_ at *ψ*_max_ = 150^°^ and *ψ*_max_ = 30^°^. The inset dark dots denotes the numerical results and the red line is the analytical solution (See ***Appendix 5***). (d,e) Coat radius *R*_coat_ vs. maximal angle *ψ*_max_ for the constant area model in (d) and for the constant curvature model in (e). Dotted lines in (d) denote the analytical solutions. Insets show the ratio of the coat radius *R*_coat_ at *ψ*_max_ = 150^°^ and *ψ*_max_ = 90^°^. The inset dark dots denotes the numerical results and the red lines are the analytical ones (See ***Appendix 5***). (f) Vesiculation diagram in the phase space of (*a*_0_, *c*_0_). Each horizontal line represents a path of the constant curvature model and each vertical line represents a path of the constant area model. Each path terminates when *ψ*_max_ = 150^°^. The solid grey lines represent contours of *ψ*_max_ = 30^°^, 60^°^, 90^°^, 120^°^, 150^°^, respectively. The solid black line is the analytical results for the vesiculation line 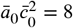(See ***Appendix 4***). The dashed black line is a random-picked straight line connecting the origin and the vesiculation boundary. The intersections of the dashed black line and the gray lines are plotted in orange dots and they are the coordinates of (*a*_0_, *c*_0_) where the five shapes in (a) are located. Shapes are arranged in an increasing order of *ψ*_max_ in (a). (b-e) Parameters with a bar over them (left vertical axes) are normalized to be dimensionless, and the dimensional parameters (right vertical axes) are calculated by one of the typical fitting values *L*_0_ = 40nm (***Figure 5***)

First, we analyze the difference between the two models in terms of membrane morphology evolution along their pathways. We fit a circle around the membrane tip and use the radius *R*_t_ of the circle to characterize the curvature of the membrane at the tip. When *R*_t_ is plotted against the maximal angle *ψ*_max_, both models show that *R*_t_ decreases with increasing *ψ*_max_ (***Figure 2***b and c). We stress that even though the intrinsic curvature *c*_0_ is fixed in the constant curvature model, it does not imply the tip radius along the vesiculation pathway is a constant. When the coating area is small, the geometric curvature at the membrane tip remains small and differs from the intrinsic curvature. Tip radius in the constant curvature model decays more steeply with *ψ*_max_ than in the constant area model. This difference becomes obvious when one plots the ratio of the tip radius at *ψ*_max_ = 150^°^ and *ψ*_max_ = 30^°^ (***Figure 2***b and c insets). For the constant area model, the ratio approximately equals to a constant 0.268 regardless of the coating area *a*_0_, which agrees well with the analytical result (See ***Appendix 5***). For the constant-curvature model, the ratio remains close to 1 only at small values of *c*_0_, as expected from the schematic representation of the model in ***Figure 1***. However, as *c*_0_ increases, the deviation from this idealized picture becomes increasingly pronounced.

Another difference between the two models is the evolution of the projected area of the clathrin coat on the substrate. We use the maximal radius *R*_coat_ of the membrane within the clathrin-coated area as the indicator of the projected area (***Figure 2***a). In the constant area model, *R*_coat_ decreases with increasing *ψ*_max_, while in the constant curvature model, *R*_coat_ increases with *ψ*_max_ and reaches a plateau (***Figure 2***d and e). The ratio *R*_coat_ (150^°^)/*R*_coat_ (90°) is about 0.732 in the constant area model and around 1 in the constant curvature model (***Figure 2***d and e, insets). The analytical curves of *R*_t_ and *R*_coat_ against *ψ*_max_ also fit perfectly with numerical solutions in the constant area model (***Figure 2***b and d, compare dotted and solid curves). Our calculations therefore demonstrate that the two models exhibit clear differences in the evolution of *R*_t_ and *R*_coat_ at the beginning of endocytosis (i.e. when *ψ*_max_ is small) which can be determined from shapes of endocytic pits obtained experimentally.

To demonstrate how the coating area *a*_0_ and the intrinsic curvature *c*_0_ of the clathrin coat influence the membrane morphology, for each pair of (*a*_0_, *c*_0_), we calculate the corresponding membrane shapes and plot the contour lines for *ψ*_max_ which indicate the stage of endocytosis (***Figure 2***f). The contour line with *ψ*_max_ = 150^°^ represents the critical line where vesiculation occurs. The line can be well fitted by the analytical expression 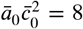 (***Figure 2***f, thick black line, see ***Appendix 4***). It implies that a small intrinsic curvature of the clathrin coat is able to induce vesiculation of the membrane with a large clathrin coat. We find that the distances between contour lines for higher values of *ψ*_max_ is smaller than those for lower values of *ψ*_max_. It means that at the late stage of endocytosis, a small change in *a*_0_ and *c*_0_ could result in a more dramatic change in the membrane shape than that at the early stage. This trend is demonstrated clearly from the orange dots in ***Figure 2***f, which correspond to shapes in ***Figure 2***a.

Note that in order to produce the phase diagram (***Figure 2***f), it requires that the bending rigidity of the clathrin coated area *κ*_coat_ is significantly larger than *κ*_bare_ in the uncoated area. If *κ*_coat_ is comparable with *κ*_bare_, there exists a region in the phase diagram in which a single (*a*_0_, *c*_0_) corresponds to three possible membrane shapes (See ***Appendix 2—Figure 1***a-d). Physically, it implies a discontinuous transition in the membrane shape along a path that passes through this region, and a gap in the maximal angle *ψ*_max_ would appear. Because in experiments, a wide spectrum of *ψ*_max_ are observed and no gap in *ψ*_max_ is found (***Avinoam et al., 2015***), we keep *κ*_coat_ much greater than *κ*_bare_ for the rest of the paper. In this regime, the membrane shapes evolve continuously along any pathway that connects the origin (0, 0) with a point on the critical vesiculation curve.

### Vesiculation needs free energy sources to drive clathrin assembly and curvature generation

In the previous section, we take curvature generation in the constant area model and clathrin assembly in the constant curvature model for granted, so that the coating area *a*_0_ and the intrinsic curvature *c*_0_ are imposed, such that the physical forces behind clathrin assembly and curvature generation are ignored. However, the bending energy *E*_b_ and the tension energy *E*_t_ typically increase with *a*_0_ and *c*_0_ along a vesiculation pathway. Therefore, vesiculation will be energetically unfavorable if no additional free energy sources are provided. In this section, we extend the model to include free energy terms for the assembly of the clathrin coat and its reorganization for curvature generation.

To describe the assembly of the clathrin coat in the constant curvature model, we introduce

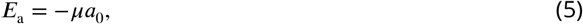

where *µ* denotes the effective surface binding energy density of clathrin molecules with the membrane. This term reduces the free energy with increasing *a*_0_, therefore, driving the assembly of clathrin. We can identify three types of free energy curves for different assembly strength *µ*: (1) When *µ* is small, the total free energy *E*_tot_ = *E*_b_ + *E*_t_ + *E*_a_ as a function of the coating area *a*_0_ has two local minima, with the lowest one at a small *a*_0_ and the other one at the maximum *a*_0_ where vesiculation occurs (***Figure 3***a, red curve). The minima are separated by an energy barrier that is significantly higher than the thermal energy *k*_B_*T* and the clathrin coat would assemble to a small area and halt. (2) With increasing *µ*, the lowest free energy minimum is shifted to the vesiculation point, but the energy barrier still exists and the clathrin coat remains small (***Figure 3***a, orange curve). (3) Vesiculation could happen for large enough *µ* such that the energy barrier vanishes and the free energy *E*_tot_ monotonically decreases with *a*_0_ (***Figure 3***a, green). Based on the above analysis of the energy landscape we construct the phase diagram of the constant curvature model with clathrin assembly in the phase space of (*c*_0_, *µ*) and classify the points into four types. Besides the three types mentioned above, when the intrinsic curvature *c*_0_ is small, increasing *a*_0_ to its maximum value (10_5_nm_2_) cannot produce vesiculation. (***Figure 3***b, gray region). The critical assembly energy density *µ* at which the energy barrier vanishes is found to increase with the intrinsic curvature *c*_0_ (***Figure 3***b, interface between the green region and the orange region), which implies that a larger assembly strength of clathrin coat *µ* is needed to complete vesiculation if the clathrin coat has a higher intrinsic curvature *c*_0_. When comparing the contour lines of the energy barrier Δ*E*_tot_ = 1*k*_B_*T* and Δ*E*_tot_ = 10*k*_B_*T* (***Figure 3***b, dotted curve and dash-dotted curve), the gap between them increases with *c*_0_, which means that the energy efficiency is reduced with *c*_0_ in the sense that, for larger *c*_0_, a larger increase in *µ* is needed to reduce the same amount of free energy.

**Figure 3.**
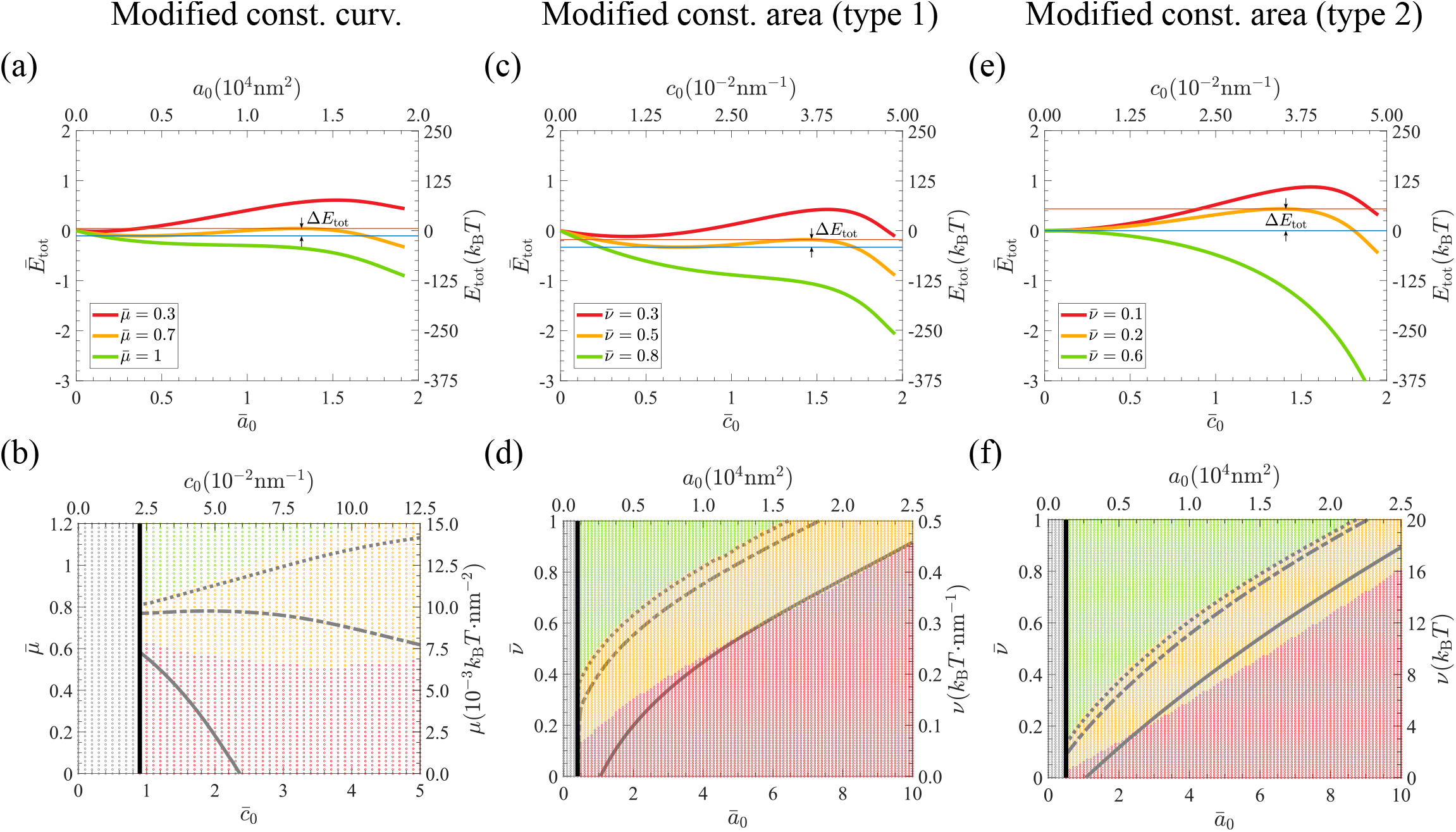
Free energy evolution in the constant curvature and constant area models when accounting for one of either the polymerization energy term *E*_a_ = −*µa*_0_ or the curvature generation energy term 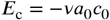 (type 1) and 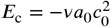(type 2). (a,b) Free energy landscape of the modified constant curvature model where 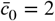 with polymerization energy *E*_a_ = −*µa*_0_ in (a) and the corresponding phase diagram in the phase space of (*c*_0_, *µ*) in (b). The rightmost endpoint of each curve is the vesiculation point where *ψ*_max_ = 150^°^. The red line in (a) and red dots in (b) correspond to pathways with minimum free energy *E*_tot_ appearing at a point other than the vesiculation point on the *E*_tot_ − *ā*_0_ curve. The orange line in (a) and the orange dots in (b) correpsond to pathways where the vesiculation point is the minimum free energy point, but an energy barrier still exists. The green line in (a) and green dots in (b) correspond to vesiculation pathways without an energy barrier. The energy barrier Δ*E*_tot_ is defined as the energy difference between the maximum point and the first minimum point before the maximum. Δ*E*_tot_ of the orange curve is shown in (a) as a typical example. The gray dots in the left-hand side of (b) correspond to pathways that numerically fail to reach the vesiculation point when *ā*_0_ reaches its upper limit 10. (c,d) Free energy landscape of the modified constant area model where *ā*_0_ = 2 with curvature generation energy *E*_c_ = −*νa*_0_*c*_0_ in (c) as a function of the intrinsic curvature *c*_0_ and the corresponding phase diagram in the phase space of (*a*_0_, *ν*) in (d). The gray dots in the left-hand side of (d) correspond to pathways that numerically fail to reach the vesiculation point when 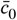 reaches its upper limit 5. (e,f) Free energy landscape of the modified constant area model where *ā*_0_ = 2 with 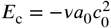 in (e) and the corresponding phase diagram in the phase space of (*a*_0_, *ν*) in (f). The gray dots in the left-hand side of (f) correspond to pathways that numerically fail to reach the vesiculation point when 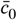 reaches its upper limit 5. (a-f) The parameters with a bar over them are normalized to be dimensionless, and the dimensional parameters are calculated using one of the typical fitting values *L*_0_ = 40nm (***Figure 5***). (b,d,f) The black line separates the region wiht gray dots (which did not numerically reach vesicultion) from the other regions. The dotted, dash-dotted and solid gray lines respectively represent the paths where Δ*E*_tot_ = 1*k*_B_*T*, Δ*E*_tot_ = 10*k*_B_*T*, Δ*E*_tot_ = 100*k*_B_*T*.

We next consider curvature generation in the constant area model. As the molecular mechanisms of curvature generation of the clathrin coat remains debated, we introduce a phenomenological model in which the free energy has the general form,

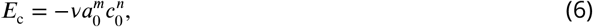

where *ν* denotes the strength of curvature generation, *m* and *n* are two positive numbers that are associated with the molecular mechanisms of curvature generation. The free energy *E*_c_ in ***Equation 6*** decreases with increasing *c*_0_, therefore, driving curvature generation. We set *m* = 1 such that *E*_c_ is proportional to the coating area. Note that *m* cannot be zero, otherwise, *E*_c_ only depends on the intrinsic curvature *c*_0_ and can be nonzero even when the coating area is zero. As for the power *n* of the intrinsic curvature *c*_0_, we set *n* = 1 or 2 (called Model(1,1) and Model(1,2), respectively). Physically, Model(1,2) implies cooperativity in the curvature generation such that the reduction of free energy per increase of unit curvature is proportional to the current curvature, i.e., Δ*E*_c_ ∝ −*c*_0_Δ*c*_0_, while in Model(1,1), the reduction of free energy per increase of unit curvature is independent of current curvature. For Model(1,1), when *ν* is small, the total free energy *E*_tot_ = *E*_b_ + *E*_t_ + *E*_c_ as a function of the intrinsic curvature *c*_0_ has two minima, the lowest one at a small positive *c*_0_ and the other one at the maximum *c*_0_ where vesiculation occurs (***Figure 3***c, red curve). Further curvature generation is strictly limited by the high energy barrier (sometimes more than 100*k*_B_*T*) between the two minima. With increasing *ν*, the lowest minimum shifts to the vesiculation point, but the energy barrier still prevents curvature generation (***Figure 3***c, orange line). For a large enough *ν*, the free energy monotonically decreases with *c*_0_ and the curvature generation proceeds until vesiculation occurs (***Figure 3***c, green curve). When the coating area *a*_0_ is very small, vesiculation fails to occur even when the intrinsic curvature is increased to its maximum value (0.125nm^−1^) (***Figure 3***c, gray region). In the phase space of (*a*_0_, *ν*), the critical value of *ν* where the energy barrier vanishes increases with the coating area *a*_0_ (***Figure 3***d, interface between the orange region and the green region), which implies that a larger clathrin coat needs a stronger strength of curvature generation to complete vesiculation.

Model(1,2) has similar free energy landscape as Model(1,1) (Compare ***Figure 3***c and e, d and f). However, in Model(1,2), for very small *ν*, the lowest free energy minimum is strictly pinned at *c*_0_ = 0, which implies no spontaneous curvature generation. In contrast, the minimum is at a small positive *c*_0_ in Model(1,1), which indicates slight curvature generation.

### Determination of the vesiculation pathway from the energy landscape

In this section, we combine the assembly energy ***Equation 5*** and the curvature generation energy ***Equation 6*** together and calculate the total free energy *E*_tot_ (*a*_0_, *c*_0_) = *E*_b_ +*E*_t_ +*E*_a_ +*E*_c_ as a function of both the coating area *a*_0_ and the intrinsic curvature *c*_0_. A pathway from the origin can be constructed by the descent along the negative gradient of the free energy landscape −∇*E*_tot_. In ***Figure 4***a we show the free energy landscape for a fixed assembly strength 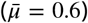 and varied reorganization strength 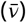 for model(1,2). When *ν* is small, the energy contour lines near the origin are kinked, which represents an energy barrier that prevents the path from going up, i.e. from generating curvature. The path extends horizontally and terminates on the *a*_0_ − *axis* (***Figure 4***a, first column, red curve). With increasing *ν*, the kinked contour lines shift towards larger *a*_0_ and the path can be lifted up to 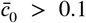 in the middle and drops to the *a*_0_ − *axis* in the end (***Figure 4***a, second column, orange curve). Beyond a critical *ν*, the energy barrier vanishes and the path bends up and terminates on the vesiculation curve (***Figure 4***a, third column, green curve). Further increasing *µ* leads to the path lifting up at a smaller coating area *a*_0_ (***Figure 4***a, fourth column, green curve). The path goes horizontally first and is later lifted up, which resembles the path of the constant area model. The three types of pathways are classified into three colored regions in the phase diagram of (*µ*, *ν*), which represent complete vesiculation (green), partial vesiculation (orange) and no vesiculation (red), respectively (***Figure 4***c).

**Figure 4.**
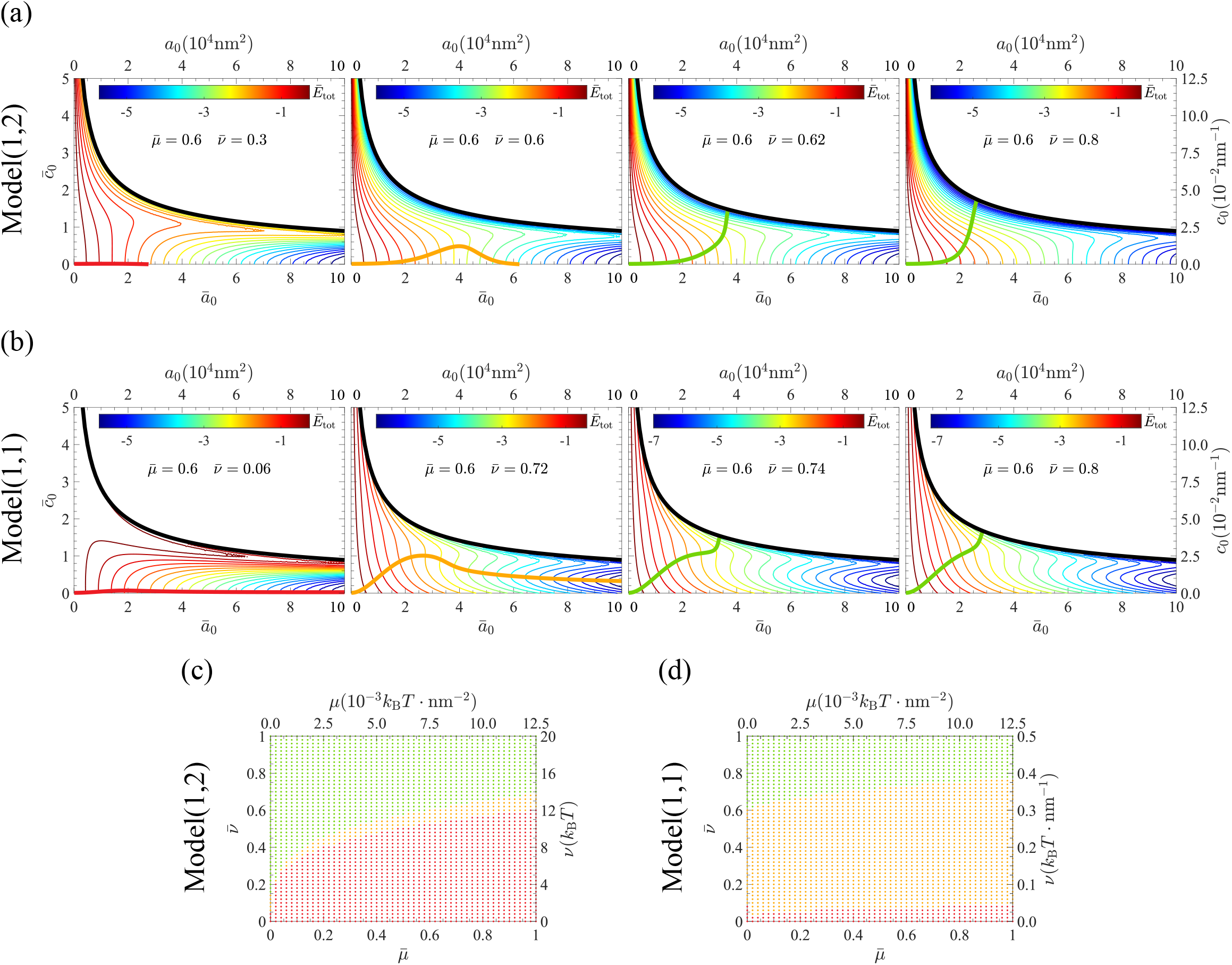
Vesiculation phase diagram when accounting for both the polymerization energy *µ* and the reorganization energy *ν*. (a,b) Free energy landscape for Model(1,2) (i.e. with reorganization energy 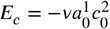) in (a) and Model(1,1) (i.e. with reorganization energy 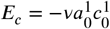) in (b). Thick black lines are the analytical solutions for the vesiculation boundary. Thin rainbow-colored lines visually represent the energy landscape (values of the color bar are for the dimensionless free energy scale). Thick colored-lines represent pathways that stream along the negative gradient of the free energy landscape in the phase space starting from the origin (i.e. no clathrin assembled and no curvatuve). Our model shows that only a subset of suitable (*µ*, *ν*) values create pathways that lead to vesiculation, i.e. that reach the thick black line (thick green lines in the third and fourth panels). The orange and red curves are pathways that fail to reach vesiculation. The red curve does not produce any curvature, while the orange curve generates a small curvature but never leads to vesiculation. (c,d) Phase diagrams for Model(1,2) in (c) and Model(1,1) in (d) show the relationship between pathway types and the (*µ*, *ν*) values. The colors of the dots correspond to the same types of pathways as represented by thick colored lines in (a,b). Our results show that larger *µ* or *ν* values lead to an easier vesiculation. Parameters with a bar over them are normalized to be dimensionless, and the dimensional parameters are calculated using one of the typical fitting values *L*_0_ = 40nm (***Figure 5***).

The free energy landscapes of Model(1,1) dramatically differs from Model(1,2). When *ν* is small, the energy gradient is strongly biased towards the horizontal direction, and the path extends horizontally with little or no curvature generation (***Figure 4***b, first column). For an intermediate *ν*, the path first goes towards the top right direction until 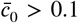 and then slowly bends down and extends towards large coating area along a valley formed in the energy landscape, which corresponds to membrane shapes with a small dimple (***Figure 4***b, second column). For large enough *ν*, the path shoots nearly straightly towards the top right direction before it reaches the vesiculation line (***Figure 4***b, third column). Further increasing *ν* makes the path more straight and terminates at a smaller coating area (***Figure 4***b, fourth column). The (*µ*, *ν*) phase diagram shows the parameter regions that lead to complete, partial or no vesiculation for Model(1,1) (***Figure 4***d).

### Comparison between different models with the experimental data

The constant area model and the constant curvature model represent two extreme pathways of membrane vesiculation. We have found constant-area-like pathways in Model(1,2) and straight-line-like pathways in Model(1,1). In order to understand which model is the most plausible, we compare membrane shapes predicted by the models with the membrane profiles obtained by electron microscopy in (***Avinoam et al., 2015***). The fitting error *ϵ* of a vesiculation path reflects the relative difference between the model-predicted geometric features along the path and the rolling median of the corresponding experimental data (***Figure 5*** and ***Appendix 3***). The fitting geometric features include neck width, tip radius, and invagination depth (***Figure 5***c). We draw the corresponding optimum energy paths that minimize the fitting error (***Figure 5***b), and compare the best model-predicted shapes with the experimental ones (***Figure 5***d).

**Figure 5.**
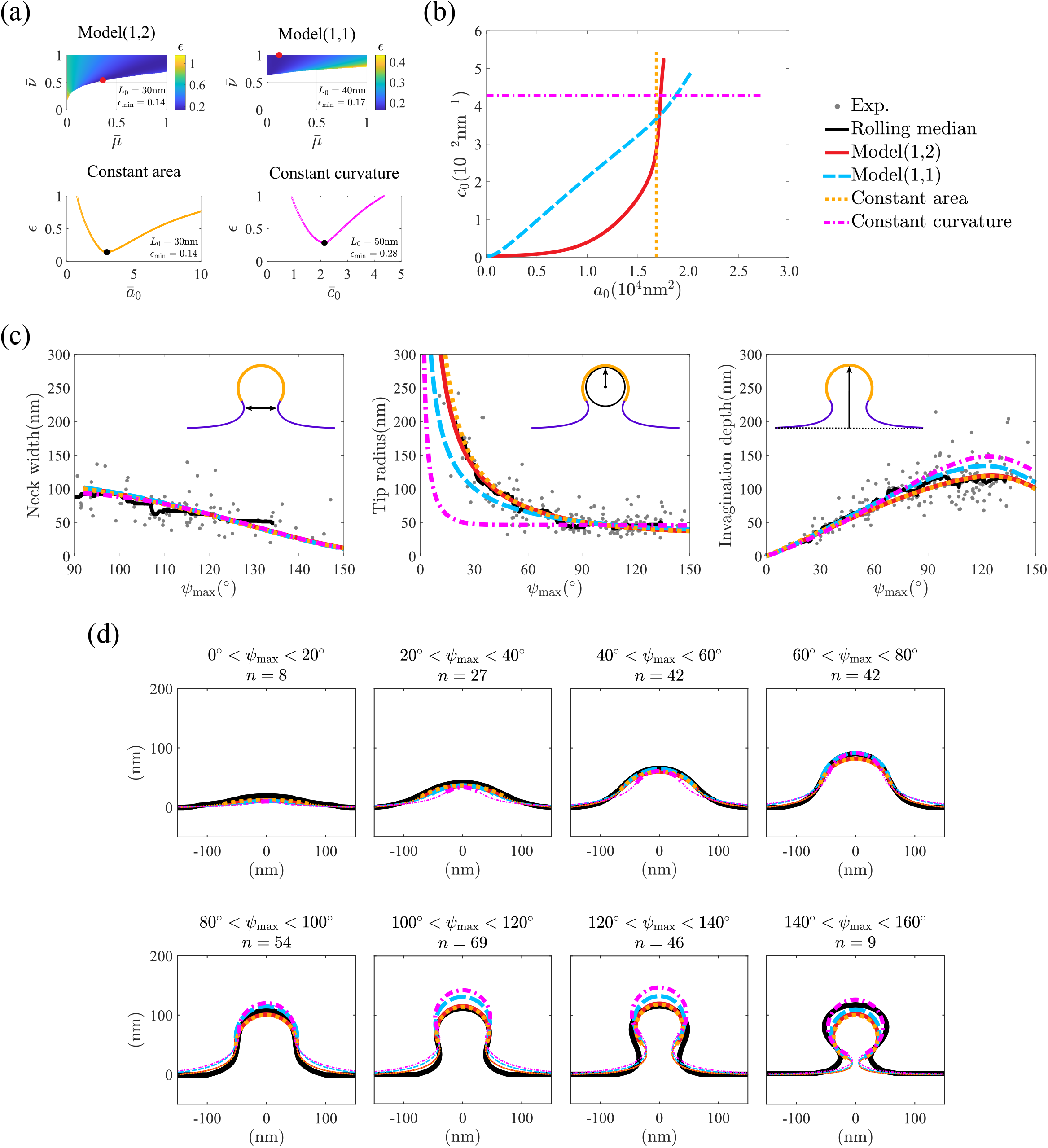
Comparison between our theory and experimental data from mammalian cells. (a) Parameter fit of the best of the four models (constant curvature model, constant area model, Model(1,1), Model(1,2)) to obtain the minimum error *ϵ*. Fitting procedure of Model(1,1) and Model(1,2) consider the total free energy *E*_tot_ = *E*_b_ + *E*_t_ + *E*_a_ + *E*_c_, while the fitting of the constant area model and the constant curvature model consider *E*_tot_ = *E*_b_ + *E*_t_. The optimized parameters are *ā*_0_ ∈ [0, 10] for the constant area model, 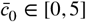 for the constant curvature model, 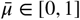 and 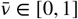 for model(*m,n*), and *L*_0_ ∈ [10nm, 100nm] within an interval of 10nm in the four models. We only assign fitting errora to the parameter sets that lead to vesiculation and only plot the error figure for the best *L*_0_. (b) Vesiculation pathways with minimum fitting error in the four models. In each model, we use the best *L*_0_ value from (a) to obtain the dimensional scale of the (*a*_0_, *c*_0_) phase space. (c) Comparison of model fits and experimental data for three geometric features: neck width, tip radius (*R*_t_) and invagination depth. Neck width is calculated as the distance between the left and right parts of the shape for *ψ*_max_ = 90^°^, and the invagination depth is measured as the height from the base to the tip of the invagination. (d) Comparison between the model-predicted shapes and the experimental shapes. Experimental membrane shapes for mammalian cells are grouped according to their maximum angle as a proxy for the different stages of CME. The number of experimental shapes falling in a certain *ψ*_max_ range is defined as *n*. The black lines are the average experimental shapes after symmetrization. The model-predicted shapes are calculated by the midpoint value of each *ψ*_max_ interval. (c,d) The curves predicted by theory are shown with colored lines, and experimental data is shown with gray dots and black lines. Parameters with a bar over them are non-dimensionalized. The detailed procedure to treat the experimental data can be found in ***Appendix 3***.

For Model(1,2) and Model(1,1), the fitting parameters include the polymerization strength 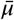 and the reorganization strength 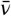 which together determine the vesiculation pathway, as well as the characteristic length *L*_0_ which scales the size of the membrane. For Model(1,2), the best fits are obtained for 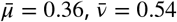 and *L*_0_ = 30nm (***Figure 5***a). The resulting path moves horizontally at first and then bents up vertically (***Figure 5***b), which resembles the behavior of the constant area model. For Model(1,1), the best fitting parameters are 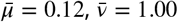 and *L*_0_ = 40nm (***Figure 5***a). The fitting path is almost a straight line towards the vesiculation line (***Figure 5***b). The optimum fitting error of Model(1,2) (*ϵ* = 0.14) is slightly better than that of Model(1,1) (*ϵ* = 0.17).

We also perform the fitting procedure to the constant area model and find the optimum parameters are *a*_0_ = 1.69 × 10^4^nm^2^ and *L*_0_ = 30nm. For the constant curvature model, the best fitting parameters are *c*_0_ = 0.043nm^−1^ and *L*_0_ = 50nm. The minimum fitting error of the constant curvature model (*ϵ* = 0.28) is exactly twice as large as that of the constant area model (*ϵ* = 0.14) (***Figure 5***a and b). So considering the fitting error and the pathway in (*a*_0_, *c*_0_) phase diagram, we raise the conclusion that the experimental vesiculation process probably favors constant-area-like pathways.

When comparing the model-predicted geometric features with the rolling median of the experimental data, we find that the four models fit almost equally well the experimental data for the neck width(***Figure 5***c left). Model(1,2) and the constant area model predict very similar results such that the curves almost overlap with each other (***Figure 5***c, red curve and orange curve). The predictions of these two models match the rolling median of the experimental data best. The constant curvature model strongly deviates from the rolling median of the experimental tip radius, particularly in the early stage of vesiculation when *ψ*_max_ < 90^°^.

To facilitate comparison between the axisymmetric membrane shapes predicted by the model and the non-axisymmetric profiles obtained from electron microscopy, we apply a symmetrization procedure to the experimental data, which consist of one-dimensional membrane profiles extracted from cross-sectional views, as detailed in Appendix 3 (see also Fig. 1). Then, we average the symmetrized profile within an interval of *ψ*_max_ ∈ [*ψ*_0_ −10^°^, *ψ*_0_ +10^°^] and overlay the averaged profile with model-predicted shapes for *ψ*_max_ = *ψ*_0_ (***Figure 5***d). At the early stage when the membrane exhibits a dimple shape (0^°^ < *ψ*_max_ < 60^°^), the membrane morphology predicted by the constant curvature model is distinguishable from the other three models, particularly when looking at the tip radius. At the late stage when the membrane exhibits an Ω-shape, i.e., *ψ*_max_ > 90^°^, the difference in shape between models is mainly manifested in the invagination depth. The constant curvature model and Model(1,1) mainly predict a deeper invagination depth than the symmetrized experimental profile, while the constant area model and Model(1,2) usually give much better fitting.

## Discussion

### Three types of clathrin coats

In this paper, we have constructed a physical model to describe how curvature generation and clathrin assembly are interrelated during the vesiculation process in CME. Previous experiments have reported three groups of clathrin coated pits, which are plaques, abortive pits, and pits that lead to vesiculation, according to their structural and dynamic properties (***Maupin and Pollard, 1983***; ***Ehrlich et al., 2004***; ***Loerke et al., 2009***; ***Saffarian and Kirchhausen, 2009***; ***Kirchhausen, 2009***; ***Lampe et al., 2016***).

In ***Figure 4***, we show that depending on the clathrin assembly strength *µ* and its reorganization strength *ν*, the pathway might end up with three possible final shapes: (i) a flat membrane with no curvature generation, (ii) a nearly flat membrane with small curvature generation, (iii) a spherical cap that leads to vesiculation. They essentially correspond to the three types of clathrin-coated pits found in experiments. Based on the phase diagram of Model(1,2) (***Figure 4***c), the difference between the three groups comes from the difference in the assembly and reorganization strengths of clathrin molecules. Furthermore, Model(1, 2) predicts that at the boundary between the type (iii) region and the type (ii) region, the reorganization strength *ν* increases with the assembly strength. This result has important implications to explain why large plaques are commonly observed in experiments. The large area of the plaques are due to the strong assembly strength *µ*. However, for these plaques to go to vesiculation, a strong reorganization energy *ν* is also needed. Therefore, the combination of a strong *µ* and weak *ν* leads to the formation of large plaques. Model(1,2) predicts that a plaque or an abortive pit can be transformed into a vesicle by either increasing the reorganization strength or reducing the assembly strength (***Figure 6***). The former ends up with a large vesicle and the latter ends up with a small vesicle. This can be used as a test of our theory with experiments to modify the binding affinity of clathrin molecules with adaptor proteins on the membrane. Weakening the affinity might increase the portion of vesicles and reduce the portion of plaques, though the vesicles would become smaller.

**Figure 6.**
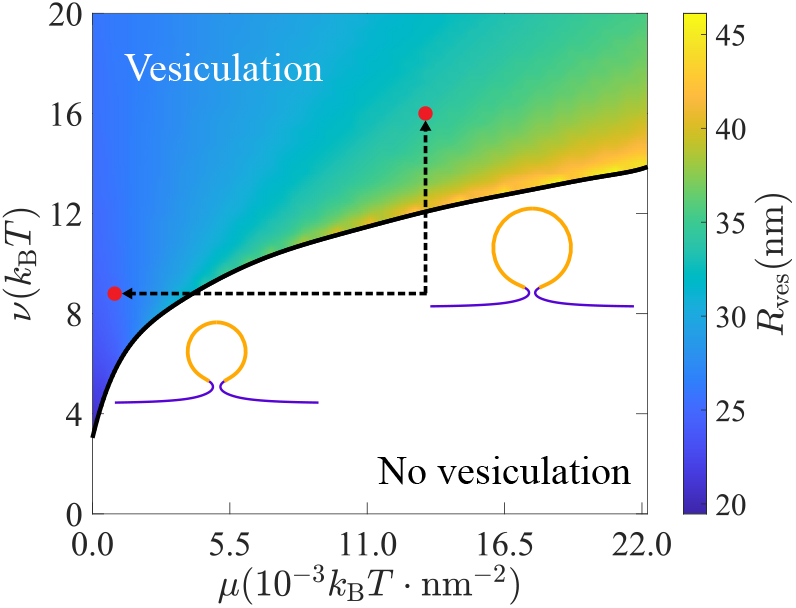
Tip radius of vesiculaion shapes (*R*_ves_) in Model(1,2). The colored region shows the (*µ*, *ν*) sets that lead to vesiculation, and brighter colors correspond to larger *R*_ves_. Decreasing assembly strength *µ* or increasing reorganization strength *ν* might lead to vesiculation of different vesicle sizes. An example from (*µ*, *ν*) = (13.3 × 10^−3^*k*_B_*T* ⋅ nm^−2^, 8.8*k*_B_*T*) to the vesiculation region is marked by arrows, red dots and corresponding vesicle shapes. The characteristic length *L*_0_ = 30nm is used in the calculation.

### Cooperativity in the curvature generation process

In ***Figure 5***, we show the best fitting results for all the four models and find that Model(1,2) produces better fits than Model(1,1), which suggests the existence of cooperativity in the curvature generation process. In particular, if curvature generation is driven by breaking bonds in the hexagonal lattice, cooperativity implies that the number of newly broken bonds is proportional to the number of already broken bonds. Because of this cooperativity, at the early stage of endocytosis bonds are broken slowly and clathrin assembly dominates over curvature generation. At the late stage of endocytosis, an increasing number of bonds are broken and curvature generation could happen rapidly and dominate over clathrin assembly. Altogether, this cooperativity leads to a constant-area-like behavior. Similar effect have been reported in Ref. (***Mund et al., 2023***).

Furthermore, although our model is purely energetic and does not explicitly incorporate dynamics, we observe in ***Figure 4***(a) that along the green curve—representing the trajectory predicted by model (1,2)—the total free energy *E*_tot_ exhibits a much sharper decrease at the late stage (near the vesiculation line) compared to the early stage (near the origin). This suggests a transition from slow to fast dynamics during endocytosis. Such a transition is consistent with experimental observations, where significantly fewer number of images with large *ψ*_max_ are captured compared to those with small *ψ*_max_ (***Mund et al., 2023***).

### Saturation effect at high membrane curvatures

Note that our model involves two distinct concepts of curvature growth. The first is the growth of *imposed* curvature—referred to here as *intrinsic curvature* and denoted by the parameter *c*_0_—which is driven by the reorganization of bonds between clathrin molecules within the coat. The second is the growth of the actual *membrane curvature*, reflected by the increasing value of *ψ*_max_. The latter process is driven by the former.

Models (1,1) and (1,2) incorporate energy terms (***Equation 6***) that promote the increase of intrinsic curvature *c*_0_, which in turn drives the membrane to adopt a more curved shape (increasing *ψ*_max_). In the absence of these energy contributions, the system faces an energy barrier separating a weakly curved membrane state (low *ψ*_max_) from a highly curved state (high *ψ*_max_). This barrier can be observed, for example, in the red curves of ***Figure 3***(a–c) and in ***Appendix 6–Figure 1***. As a result, membrane bending cannot proceed spontaneously and requires additional energy input from clathrin assembly.

The energy terms described in ***Equation 6*** serve to eliminate this energy barrier by lowering the energy difference between the uphill and downhill regions of the energy landscape. However, these same terms also steepen the downhill slope, which may lead to overly aggressive curvature growth. To mitigate this effect, one could introduce a saturation-like energy term of the form

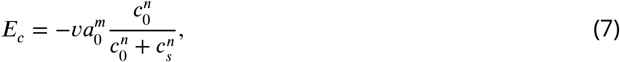

where *c*_*s*_ represents a saturation curvature. Importantly, adding such a term would not alter the conclusions of our study, since the energy landscape already favors high membrane curvature (i.e., it is downward sloping) even without the additional energy terms.

### The difference in membrane morphology between the different models is most salient at the early stage of endocytosis

When we compare the model predictions, we find that the difference in membrane morphology between models is not as big as expected, which might explain why it has been difficult to distinguish between the constant area and the constant curvature models for so long. For instance, the neck width vs. *ψ*_max_ and the invagination depth vs. *ψ*_max_ are similar for all the four models (***Figure 5***c, left and right). The best fitting error of the four models calculated from the geometric features are relatively close, except for the constant curvature model, which gives the worst fit (***Figure 5***a). The models are mainly distinguishable from the tip radius vs. *ψ*_max_ plot at the early stage of endocytosis when the membrane is nearly flat (***Figure 5***c, middle), i.e. for shapes with small *ψ*_max_. However, published experimental shape at early stages of endocytosis are sparse. Our result hints that in order to distinguish between the models, collecting membrane shapes at the early stage is necessary and the relation of tip radius vs. *ψ*_max_ is the key geometric feature to tell the models apart.

### The projected area of clathrin coat in the plane of the plasma membrane could distinguish between the two models

In ***Figure 2***d and e we have shown that the coat radius *R*_coat_, which represents the projected area of the clathrin coat in the plane of the plasma membrane, as a function of *ψ*_max_ exhibit opposite trends for the constant curvature model and the constant area model. The results suggest that in experiments the projected area for the constant area model would first increase and then decrease over time, finally reaching a plateau. However, for the constant curvature model, the projected area would increase over time and finally reach a plateau without a decreasing phase. This result suggests another method to distinguish between the two models via the projected area measurement. The idea has been used in a study where the clathrin-coated pit was imaged with platinum replica and cryoelectron microscopy and tomography (***Sochacki et al., 2021***). The results support a constant-area-like model, consistent with the prediction of our Model(1,2), in which the dome structures have a slightly larger coating area than flat structures. On the other hand, another study has used the super-resolved live cell fluorescence imaging with TIRF to measure the growth of the clathrin coat over time (***Willy et al., 2021***). The authors found a smooth drop in the projected area of clathrin coat over time. However, based on a computer simulation of clathrin assembly, they concluded that the smooth drop of the projected area was the result of a constant-curvature-like model because a constant-area-like model would produce a sharp drop. We attribute the difference between their model and our model to the fact that they model the clathrin coat as a discrete lattice while we use a continuum mechanics method. More importantly, in their model, the moment at which curvature generation occurs was arbitrarily imposed at 80% of clathrin triskelions. If the transition were chosen to occur with fewer triskelions, e.g. 40%, the sharp drop in the project area might not happen in the constant-area-like model. Furthermore, the authors used the number of triskelions to monitor the progress of endocytosis which terminates when the triskelions reach the maximum number. This choice might bias towards the constant-curvature-like model because the vesiculation may not happen at all when the clathrin numbers reaches its maximum.

**Table 1.** List of default parameters in the model.

| Symbols | Meaning | Value | Unit |
| --- | --- | --- | --- |
| $\kappa_{\text{bare}}$ | Bending rigidity of uncoated area | 20 | $k_B T$ |
| $\kappa_{\text{coat}}$ | Bending rigidity of coated area | 1000 | $k_B T$ |
| $\sigma_e$ | Membrane tension at the base | 0.025 | $\text{pN} \cdot \text{nm}^{-1}$ |

### The bending rigidity of the coated area should be much larger than the uncoated area

Comparison of our model to experimental data demonstrates that the relative bending rigidities of the coat and the membrane are constrained. Indeed, if *κ*_coat_ /*κ*_bare_ < 50, the model predicts an abrupt change (or a gap) in *ψ*_max_ at the end of vesiculation (See ***Appendix 2—Figure 1***), i.e., a snap-through transition (***Hassinger et al., 2017***). If a gap in *ψ*_max_ existed, we would expect that the distribution of experimental shapes to be discontinuous, with no or very few data corresponding to a certain range of *ψ*. However, in the experiments (***Avinoam et al., 2015***), the endocytic pits shapes are continuously distributed across the *ψ*_max_ spectrum, indicating the ergodicity of *ψ*_max_ during the endocytic process. Our calculation suggests that the clathrin coat is about 50 times stiffer than the membrane.

### Comparison with particle wrapping

The purpose of the clathrin-mediated endocytosis studied in our work is the recycling of membrane and membrane-protein, and the cellular uptake of small molecules from the environment — molecules that are sufficiently small to bind to the membrane or be encapsulated within a vesicle. In contrast, the uptake of larger particles typically involves membrane wrapping driven by adhesion between the membrane and the particle, a process that has also been studied previously (***Góźdź, 2007***; ***Bahrami et al., 2016***). In our model, membrane bending is driven by clathrin assembly, which induces curvature. In particle wrapping, by comparison, the driving force is the adhesion between the membrane and a rigid particle. In the absence of adhesion, wrapping increases both bending and tension energies, creating an energy barrier that separates the flat membrane state from the fully wrapped state. This barrier can hinder complete wrapping, resulting in partial or no engulfment of the particle. Only when the adhesion energy is sufficiently strong can the process proceed to full wrapping. In this context, adhesion plays a role analogous to curvature generation in our model, as both serve to overcome the energy barrier. If the particle is spherical, it imposes a constant-curvature pathway during wrapping. However, the role of clathrin molecules in this process remains unclear and will be the subject of future investigation.

## Acknowledgments

We thank Prof. Ori Avinoam and Marko Kaksonen for generously sharing their data with us. R.M. acknowledges financial support from Fundamental Research Funds for Central Universities of China under Grant No. 20720240144. Part of this work was funded by NIH R01 grant GM115636 awarded to J.B.

## Appendix 1

### Detailed formula derivation

We assume the membrane shape is axisymmetric and is parameterized with its meridional curve {*r*(*u*), *z*(*u*)}, with *u* = 0 corresponding to the membrane tip and *u* = 1 corresponding to the membrane edge. Our aim is to derive the shape equations via minimizing the bending energy of the membrane under certain geometric constraints (***Jülicher and Seifert, 1994***; ***Seifert et al., 1991***; ***Zhong-Can and Helfrich, 1987***).

Hereafter we use *f*^′^ ≡ d*f*/d*u* to denote the derivative of an arbitrary function *f* with respect to the rescaled arclength *u*. It should be noticed that if {*r*(*u*), *z*(*u*)} describes a membrane shape, {*r*[*g*(*u*)], *z*[*g*(*u*)]} describes the same membrane shape if the function *g* maps the interval [0, 1] to [0, 1], e.g., *g*(*u*) = *u*^2^. In order to fix the issue, we introduce 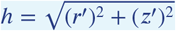 and impose that *h* is a constant. By this definition, we essentially let *u* = *s*/*S*, where *s* is the arclength calculated from the membrane tip and *S* is the total arclength. The constant *h* = *S* is an unknown parameter to be determined via solving the shape equations. In order to simplify the form of the bending energy, we introduce the tangent angle *ψ*(*u*) which satisfies the geometric relation:

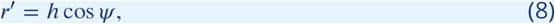

and

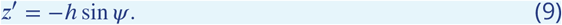

The two principal curvatures can be expressed as

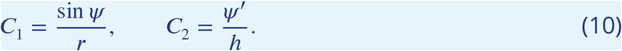

The bending energy of the membrane then reads

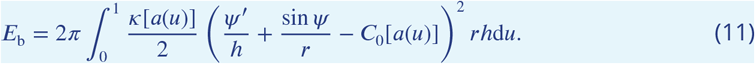

Note that the bending rigidity *κ*[*a*(*u*)] and the intrinsic curvature *C*_0_[*a*(*u*)] are functions of the area *a*(*u*), which fullfils the equation

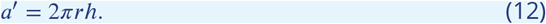

This relationship means that we study an inhomogeneous membrane which is locally incompressible in its area. In order to impose the geometric relation Eqs. (8),(9) and (12), we introduce three Lagrangian multipliers *γ*(*u*), *η*(*u*) and *σ*(*u*) to the integral

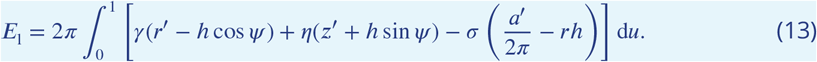

The total free energy to be minimized under the geometric constraints reads

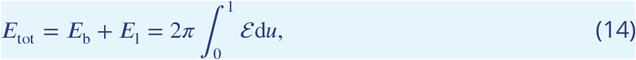

in which ℰ reads

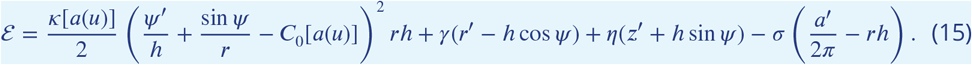

The variation of the functional *E*_tot_ in ***Equation 14*** reads

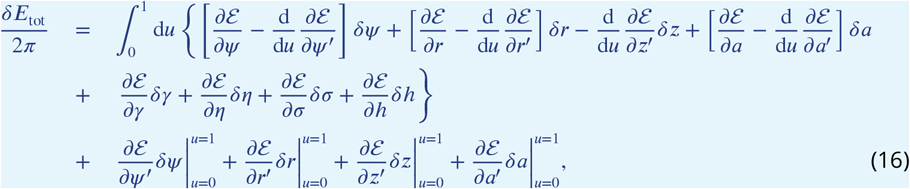

which contains both the bulk terms (first line) and the boundary terms (second line). The Euler-Lagrange equations can be obtained by the vanishing of the former 7 bulk terms, which are reduced to

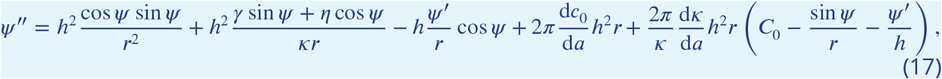

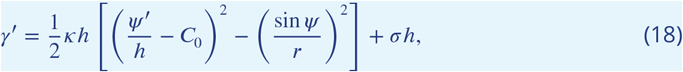

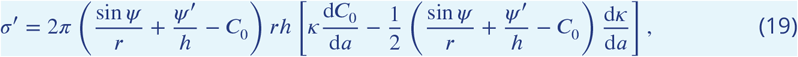

and

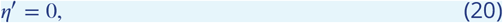

as well as Eqs. (8), (9) and (12). Note that the vanishing bulk term of *δh* gives us

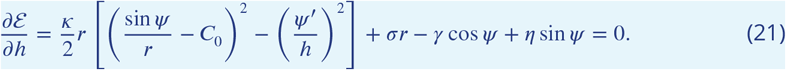

which is a boundary condition (not one of the differential equations).

Next, we need 9 boundary conditions (BCs) to numerically solve 6 first-order differential equations, 1 second-order differential equation together with 1 unknown parameter. At the membrane tip *u* = 0, we can easily obtain *r*(0) = 0, *ψ*(0) = 0, *a*(0) = 0 by geometric relations. Then, *∂* ℰ /*∂h* = 0 given by ***Equation 21*** is satisfied at *u* = 0. At the membrane base *u* = 1, we can also easily get *r*(1) = *R*_b_, *z*(1) = 0 by geometric relations, where *R*_b_ is the distance from the boundary *u* = 1 to the axisymmetric axis. All boundary terms in ***Equation 16*** vanish simultaneously because geometric relations lead to *δf* = 0, except *δz* at *u* = 0 and *δψ* at *u* = 1. This forces *∂* ℰ /*∂z*^′^ = 0, which is, *η*(0) = 0, and *∂* ℰ /*∂ψ*^′^ = 0, and leads to

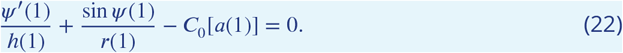

Surface tension is fixed to *σ* = *σ*_e_ at the base. The definition of the characteristic length is 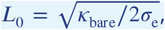 depicting the radius of a tube of membrane elongated by a point force.

In summary, all the BCs are listed below

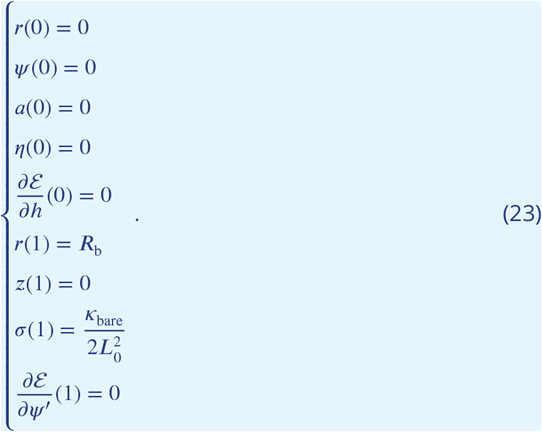

We adopt 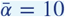 and 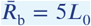 in all of our simulation. Other more important variables are already given in the main text.

## Appendix 2

### Gap of maximum angle

All of our results in the main text were computed under the assumption of a very large ratio *κ*_coat_ /*κ*_bare_ = 50 to prevent a discontinuous jump of *ψ*_max_. When *κ*_coat_ /*κ*_bare_ = 5, the gap of *ψ*_max_ is observable in some (*a*_0_, *c*_0_) sets (***Appendix 2—Figure 1***). For the constant area model, the gap is observed around *ψ*_max_ = 125^°^ when *a*_0_ is above a limit value and increases slightly with increasing *a*_0_ (***Appendix 2—Figure 1***a). The solid and dotted lines deviate further with an increasing *a*_0_. The dotted free energy curve with larger *a*_0_ generates a Gibbs triangle (***Appendix 2—Figure 1***c). The bottom point of the Gibbs triangle is the phase change point and corresponds to the *c*_0_ value where the gap of *ψ*_max_ is situated on. When *a*_0_ is less than the limit value, the free energy curve is smooth and has no obvious phase change point. Result in the constant curvature model is qualitatively similar to the constant area model (***Appendix 2—Figure 1***b and d). If *ψ*_max_ gap exists, the dotted line and solid line intersect at three multiple-solution points with the same *c*_0_ and *a*_0_.

The (*a*_0_, *c*_0_) phase diagram shows how both arguments effect on *ψ*_max_ gap (***Appendix 2—Figure 1***e). The orange lines shows the deviation and variation trend between the dotted lines and the solid lines in ***Appendix 2—Figure 1***a and b. Note that the deviation of any path that starts from the origin and terminates on the vesiculation boundary can be described by orange lines, rather than just constant curvature paths or constant area paths. A physical vesiculation pathway just goes across three orange lines directly and terminates at the vesiculation boundary while a numerical one turns back at the upper line, then turns back at the lower line, and finally completes vesiculation. Note that some section of the upper dotted curve overlaps with the numerical vesiculation boundary, because the system reaches vesiculation boundary before passing through the phase change point. Finally, the introduction of the assembly energy and the reorganization energy have no effect on *ψ*_max_ gap, and just change the shape of the Gibbs triangle.

### Boundary radius value

We set 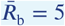 throughout our study. Values of tip radius and neck radius appear almost identical for different *R*_b_ in both models (***Appendix 2—Figure 2***). However, membrane heights have distinct differences for different *R*_b_ when *ψ*_max_ is in the middle range. This difference is likely due to fact the uncoated area contains a smaller region able to generate curvature and does not contribute in lifting the membrane center at this stage. From 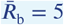 to 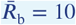, the curves are virtually identical, even for membrane height. Since a larger *R*_b_ requires a larger number of mesh points in the simulations, we chose 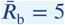 to balance computation time and numerical accuracy.

**Appendix 2—figure 1.**
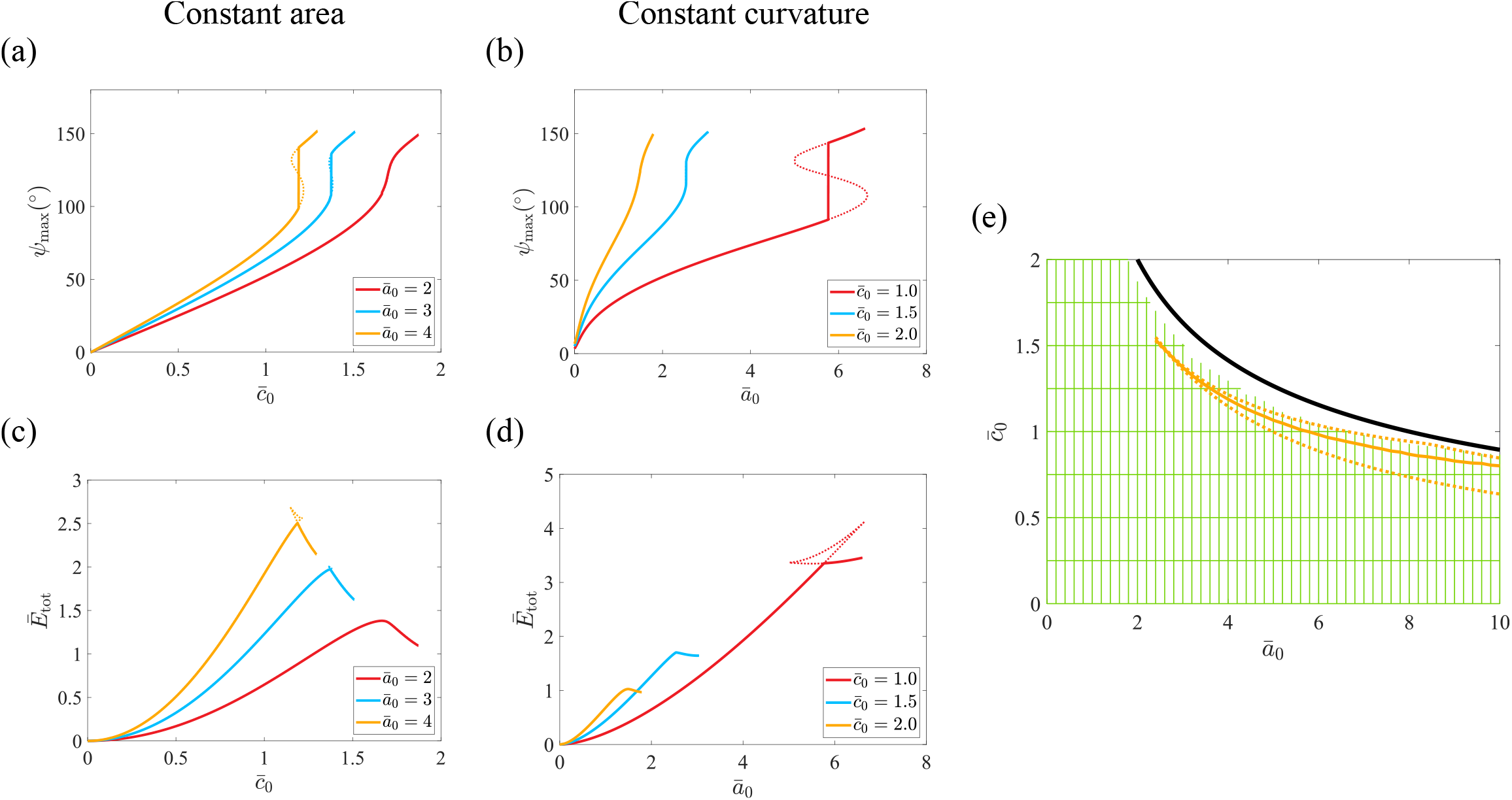
Vesiculation pathways and free energies in the different models studied in this paper. (a,b) *ψ*_max_ for the constant area model and the constant curvature model. (c,d) Free energy for the constant area model and the constant curvature model. (a-d) Colored lines represent vesiculation pathways for a fixed *a*_0_ or *c*_0_ specified in the legend. Solid lines represent states that have the minimum free energy, while dotted lines represent energetically possible states but are metastable. For certain range of *a*_0_ or *c*_0_, a single of *a*_0_ or *c*_0_ corresponds to multiple values of *ψ*_max_. In the free energy diagram, this is reflected in the Gibbs triangle, which means there will be a snap-through transition of *ψ*_max_. (e) Vesiculation boundary in the (*a*_0_, *c*_0_) phase diagram. Each green line represents a pathway in one of the two models (horizontal lines for the constant curvature model, vertical lines for the constant area model). Each line stops when vesiculation occurs (i.e. *ψ*_max_ = 150^°^) according to the numerical simulations of the model. The black line represents the vesiculation boundary as determined analytically, i.e. when 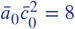 (See ***Appendix 4***). The solid orange line represents (*a*_0_, *c*_0_) values for which a *ψ*_max_ gap exists. The region between the two dotted lines represent that a single pair of (*a*_0_, *c*_0_) corresponds to three values of *ψ*_max_, and the solid line represents the critical line at which a snap-through transition of the shape will occur. Note that the vesiculation boundaries detemined numerically or analyticaly poorly match to each other because the ratio *κ*_bare_/*κ*_coat_ is small and ***Equation 31*** is not satisfied (See ***Appendix 4***).

**Appendix 2—figure 2.**
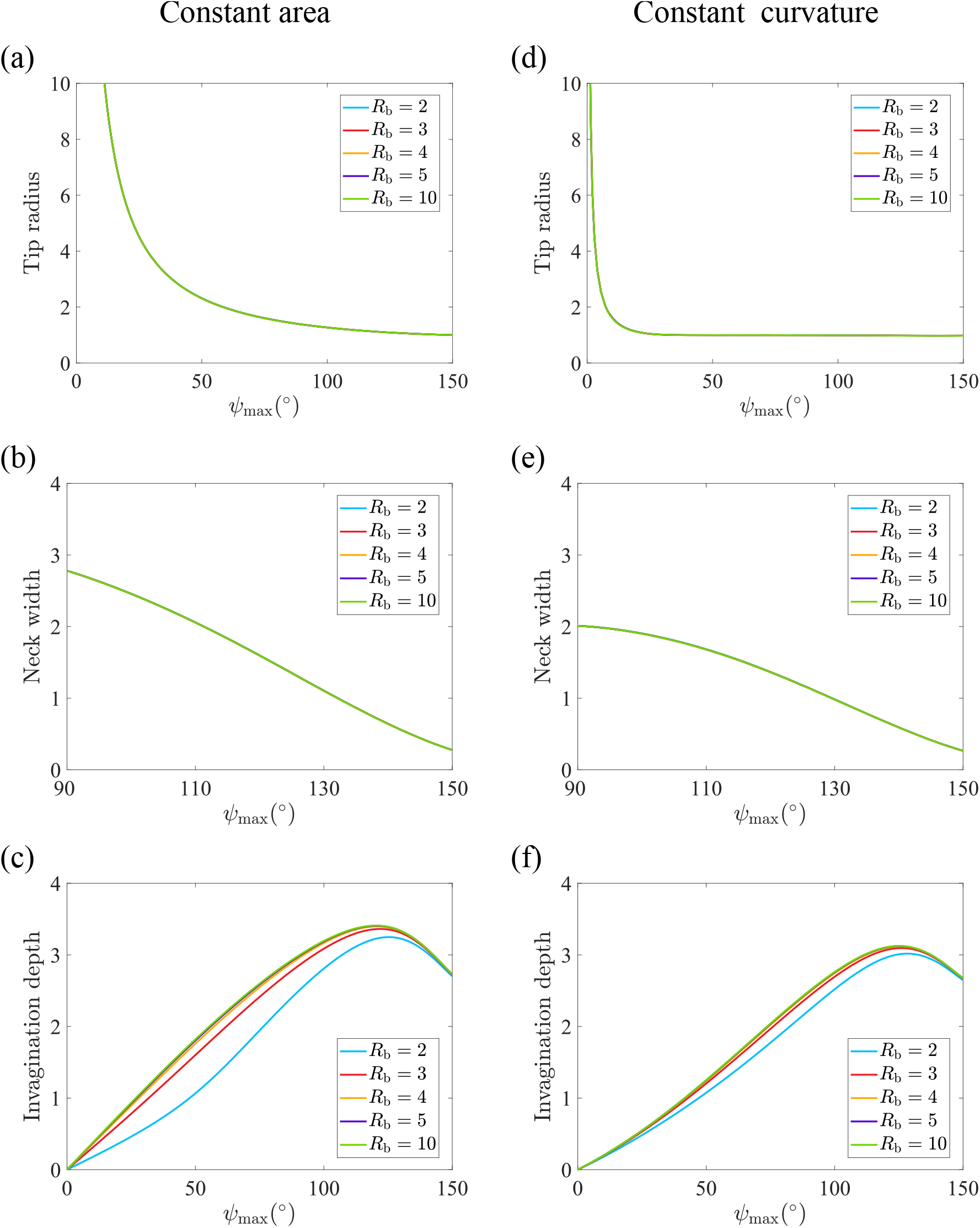
Influence of *R*_b_ on the shape parameters in the constant area and constant curvature models. (a and d) Neck width. (b and e) Tip radius (note that the neck width is ill-defined when *ψ*_max_ is small, so we restrict the abscissa range, and the membrane height is *z*(*u*) at *u* = 1). (c and f) Membrane height. (a-c) Constant area model. (d-f) Constant curvature model. Default parameters used in this figure: 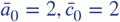.

## Appendix 3

### Symmetrization algorithm

In our investigation to determine the model parameters from experimental data, we need to symmetrize the shapes extracted from electron tomograms using the paired coordinates (*r*_*i*_, *z*_*i*_)(***Appendix 3—Figure 1***a is an example). Firstly, we define the vertical line that crosses the maximum *z*-value of the shape as its axisymmetric axis, and normalize the left- and right-poritons of shapes using the same rescaled mesh points *u* = *s*_*i*_/*S*_*i*_. Secondly, we average *r* and *z* with the same *u* on both sides to achieve the symmetrized shape (***Appendix 3—Figure 1***b).

### Rolling median calculation

We note (*x*_1_, *y*_1_), (*x*_2_, *y*_2_), …, (*x*_*i*_, *y*_*i*_), …, (*x*_*N*_, *y*_*N*_) the coordinates of all *N* data points in a given figure (Fig. 5c), sorted by independent variable *x*_*i*_ in an ascending order. We calculate the median points 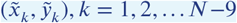 of ten consecutive neighbouring points (*x*_*i*_, *y*_*i*_), …, (*x*_*i*+9_, *y*_*i*+9_) by *x* and *y*, respectively. We connect consecutive median points to plot the rolling median line. The same procedure is performed for each of the three parameter figures.

The symmetrized experimental shapes are grouped by their *ψ*_max_ value in eight intervals of equal width ranging from *ψ*_max_ = 0^°^ to *ψ*_max_ = 160^°^ (***Figure 5***d). A single range of *ψ*_max_ contains *n* shapes {*r, z*}_1_, …, {*r, z*}_*n*_. The {*u*}_*n*_ meshes being exactly the same for each *n* (See previous section), we calculate the average values in each *ψ*_max_ interval as

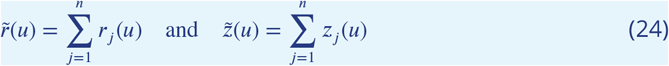

to obtain eight average shapes 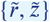. Then, we draw the shapes of four theoretical models with the midpoint *ψ*_max_ values for each interval to compare our theoretical shapes with the averaged experimental shapes.

### Error calculation

We compare four models with the rolling median lines, and draw the relationship between total relative fitting error and parameters (***Figure 5***a and c). In the *ψ*_max_ interval where theoretical and experimental data points are well-defined, we use an interpolation method to map their *ψ*_max_ into the same mesh points. The total relative fitting error is calculated by

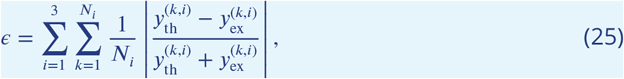

where *N*_*i*_ is the number of well-defined experimental data points in the *i*-th parameter figure, and 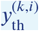 is the *k*-th theoretical value in the *i*-th para figure, and 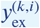 is the *k*-th experimental value in the *i*-th para figure. We choose the parameter sets with minimum *ϵ* in four theory models. Then, we compare the curves and shapes obtained using the best-fit parameters with the experimental data. We only show the error-parameter figure with the best-matched *L*_0_ value in Fig. 5a.

### Normalization

We impose *Ā* as the dimensionless variable of *A* and *A*_unit_ as the normalizing units, such that *A*/*A*_unit_ = *Ā*. Units of model variables are listed in ***Appendix 3—Table 1***. Other variables are derived variables and are not listed in the table.

**Appendix 3—figure 1.**
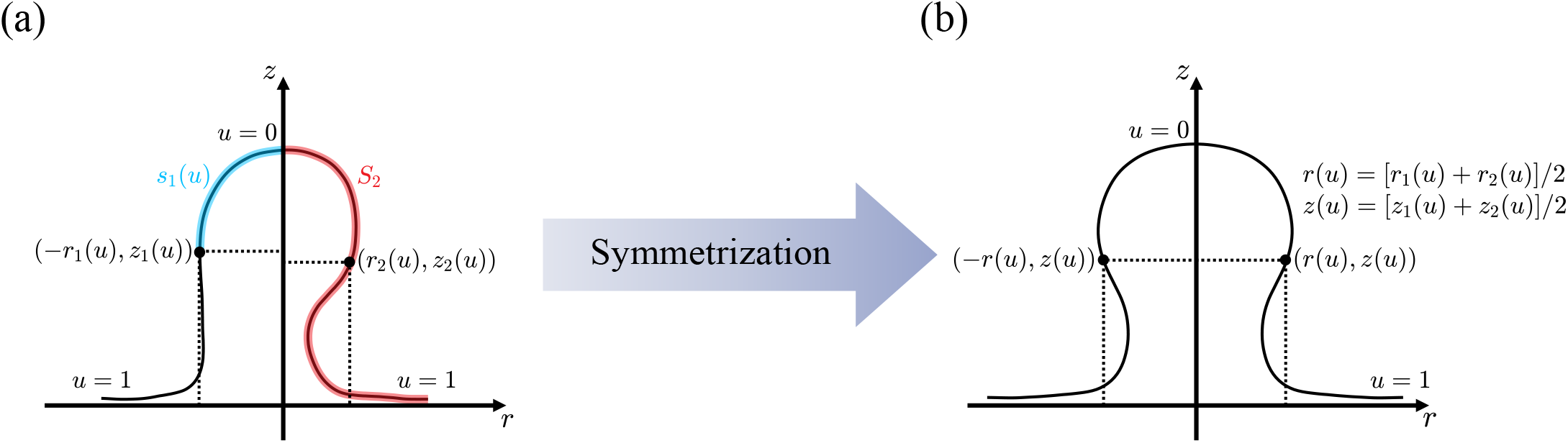
Symmetrization of the experimental profile. (a) The experimental profile curve is divided into a left part and a right part using the highest point to split the profile. Each part is interpolated onto the same rescaled mesh points *u*_*i*_ = *s*_*i*_/*S*_*i*_, where *s*_*i*_ is the arclength calculated from the highest point, *S*_*i*_ is the total arclength to the last point and *i* = 1, 2 indicates the left or right section, repectively. The length of the translucent blue line is *s*_1_, and the translucent red line is *S*_2_. (b) Symmetrized experimental profile by taking the average of the left part and the right part at the same rescaled arclength.

**Appendix 3—table 1.** Model parameters and their units.

| Parameters | Meaning | Unit |
| --- | --- | --- |
| $u$ | Independent variable with fixed boundary. | Dimensionless. |
| $\psi$ | Tangent angle of the membrane. | Dimensionless |
| $m, n$ | Powers in Model($m, n$). | Dimensionless. |
| $L_0$ | Characteristic length. | 40nm |
| $h$ | Assistant variable. | $L_0$ |
| $r, z$ | Geometric coordinates. | $L_0$ |
| $s$ | Arc length. | $L_0$ |
| $S$ | Total arc length. | $L_0$ |
| $R_b$ | Base radius. | $L_0$ |
| $C_1, C_2$ | Principal curvatures of the membrane. | $L_0^{-1}$ |
| $C_0$ | Intrinsic curvature of a certain point on the membrane. | $L_0^{-1}$ |
| $c_0$ | Intrinsic curvature of the clathrin. | $L_0^{-1}$ |
| $a$ | Area. | $2\pi L_0^2$ |
| $a_0$ | Coating area of clathrin. | $2\pi L_0^2$ |
| $A$ | Total area of the membrane. | $2\pi L_0^2$ |
| $\alpha$ | Sharpness of the step function | $(2\pi L_0^2)^{-1}$ |
| $\kappa$ | Bending rigidity of a certain point on the membrane. | $20k_B T$ |
| $\kappa_{\text{bare}}$ | Bending rigidity of bare membrane. | $20k_B T$ |
| $\kappa_{\text{coat}}$ | Bending rigidity of coated membrane. | $20k_B T$ |
| $\mathcal{E}$ | Differential energy. | $20k_B T$ |
| $E_b$ | Bending energy. | $2\pi \cdot 20k_B T$ |
| $E_t$ | Tension energy. | $2\pi \cdot 20k_B T$ |
| $E_a$ | Area assembly energy. | $2\pi \cdot 20k_B T$ |
| $E_c$ | Curvature generation energy. | $2\pi \cdot 20k_B T$ |
| $E_l$ | Lagrangian energy term. | $2\pi \cdot 20k_B T$ |
| $E_{\text{tot}}$ | Total free energy. | $2\pi \cdot 20k_B T$ |
| $\gamma, \eta, \sigma$ | Lagrangian multipliers. | $20k_B T \cdot L_0^{-1}$ or $20k_B T \cdot L_0^{-2}$ |
| $\sigma_e$ | Membrane tension at the base. | $20k_B T \cdot L_0^{-2}$ |
| $\mu$ | Assembly strength. | $20k_B T \cdot L_0^{-2}$ |
| $\nu$ | Reorganization strength. | $2 \cdot 40^{1-2m+n} \cdot 20k_B T \cdot L_0^{-2+2m-n}$ |

## Appendix 4

### Critical vesiculation curve

We approximate the system as a combination of a spherical cap and a plane (***Appendix 4—Figure 1***, cross-section view). The undeformed shape of the coating area is a spherical cap with curvature *c*_0_ and the uncoated area is flat. For simplicity, We postulate the spherical cap deforms uniformly and the plane has no deformation. From the geometric relationship, the area can be written as

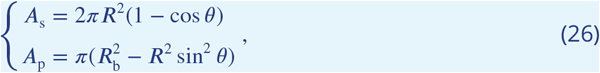

where *A*_s_ is the area of spherical cap and *A*_p_ is the area of plane. Correspondingly, their surface bending energy density expressions are

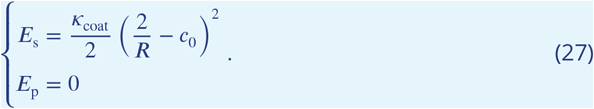

The uncoated area has no deformation so it has no bending energy. The total free energy includes the surface tension, so

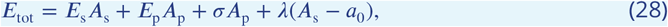

where *λ* is the Lagrange multiplier to give geometric restriction on *A*_s_ = *a*_0_, and *σ* = *σ*_e_ is a constant surface tension of uncoated area. Physically, the system prefers the shape that minimizes *E*_tot_, i.e.

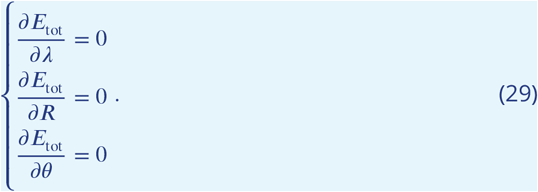

Eliminating *λ* and *R* and ignoring solutions where *θ* < 0, which are not physiclaly possible, and *θ* = *π*, which can only be achieved after passing an energy barrier, we obtain

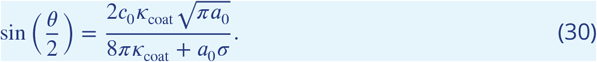

Then, we postulate that the rigidity of the solid shell is very large, giving 8*πκ*_coat_ ≫ *a*_0_*σ*. We can then use the approximation:

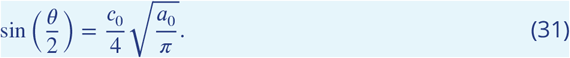

Then, we consider a closed sphere as the vesiculation state, where *θ* = *π*. Finally, we substitute 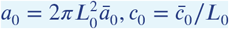, to obtain the dimensionless relation

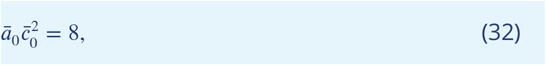

which is the analytical vesiculation boundary of this simplified model.

## Appendix 5

### Model fitting

Using the approximated model (***Appendix 4—Figure 1***), we provide an analytical result to distinguish between the constant area model and the constant curvature model (***Figure 2***b-e). This result is confirmed by numerical analyses and can be used to differentiate the two models from experimental data. The area of a spherical cap is

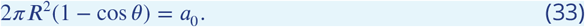

In the constant area model, *R* and *θ* vary during the vesiculation process but *a*_0_ remains contant. ***Equation 33*** gives

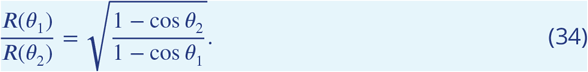

Therefore, we calculate the ratios *R*_t_ (150^°^)/*R*_t_ (30^°^) ≈ 0.268 and *R*_coat_ (150^°^)/*R*_coat_ (90^°^) ≈ 0.732, which both are close to the results from the numerical simulations. In the constant curvature model, *θ* and *a*_0_ vary during the vesiculation process but *R* remains constant, so

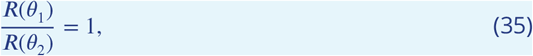

which holds true for any *θ*. So, in the analytic model, *R*_t_ (150^°^)/*R*_t_ (30^°^) = 1 and *R*_coat_ (150^°^)/*R*_coat_ (90^°^) = 1. The latter ratio matches the numerical results well but the former ratio doesn’t because of the very large tip deformation that significantly differ from a spherical cap in some shapes determined numerically. Using different pairs of *ψ*_max_ values, *R*_t_ and *R*_coat_ ratios are different in both models, and these ratios can be used as indicative variables to distinguish between the two models from the experimental shapes.

In addition, the simplified model postulates *R* = *R*_t_, *θ* = *ψ*_max_ and ***Equation 33*** leads to

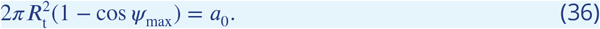

For *R*_coat_ the analytical expressions are

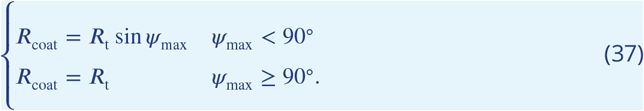

***Equation 36*** and ***Equation 37*** both fits the numerical results well (***Figure 2***b-e), proving the validity of the spherical cap approximation.

**Appendix 5—figure 1.**
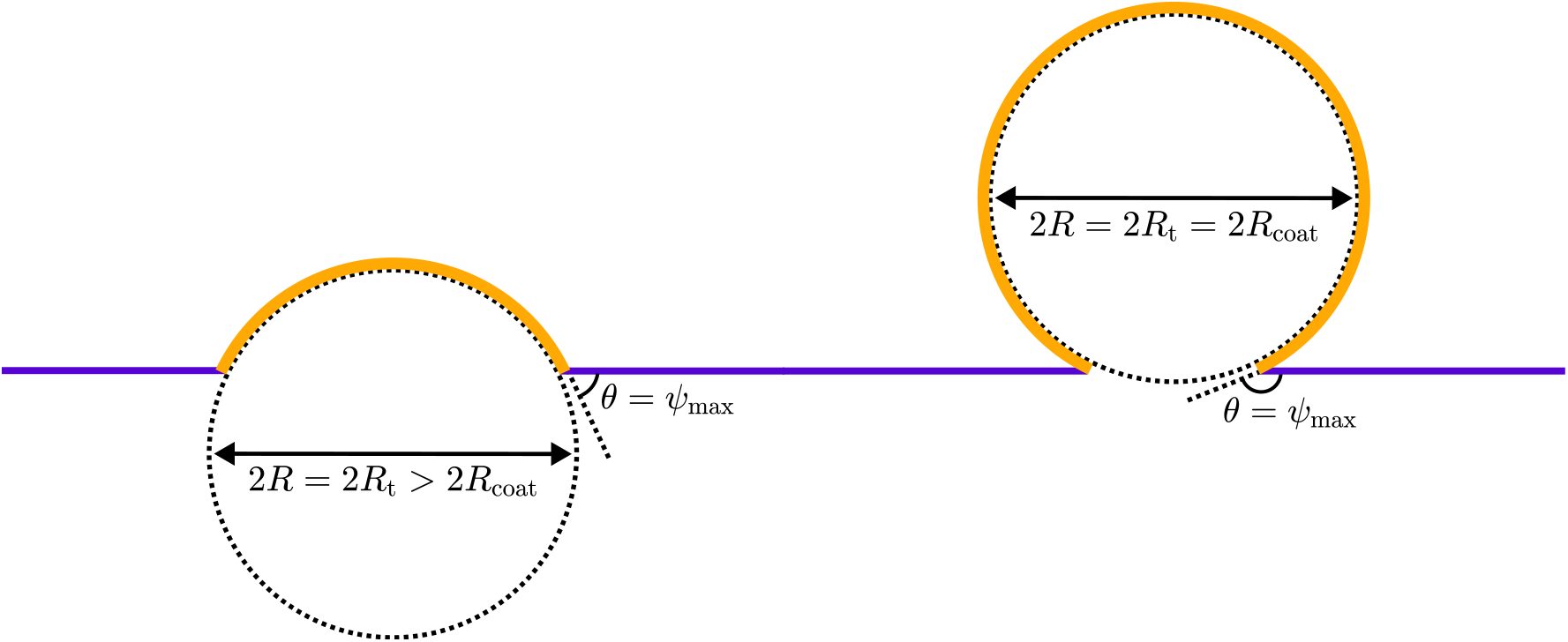
Approximate spherical cap model at small *ψ*_max_ (left) and large *ψ*_max_ (right). The radius of the spherical cap is *R*, the max tangential angle of the sphere is *ψ*_max_, equaling to the angle at the base. Other definitions are the same as in the main text. *R* = *R*_t_ > *R*_coat_, *θ* = *ψ*_max_ when *ψ*_max_ < 90^°^ and *R* = *R*_t_ = *R*_coat_, *θ* = *ψ*_max_ when *ψ*_max_ ≥ 90^°^.

## Appendix 6

### Decomposition of membrane energy

The total membrane energy consists of bending energy *E*_b_ and tension energy *E*_t_. In ***Appendix 6–Figure 1***, we present *E*_b_ and *E*_t_ as functions of the coating area a_0_ and the intrinsic curvature *c*_0_, as a supplement to the main energy landscape shown in ***Figure 4***. Our analysis reveals that tension energy dominates over bending energy, particularly at large coating areas *a*_0_. For both energy components, an energy barrier separates the slightly bent mem brane state (low *ψ*_max_) from the vesiculation boundary (high *ψ*_max_). The inclusion of assembly energy *E*_a_ and reorganization energy *E*_c_ is intended to eliminate this barrier. Since the tension energy scales with the membrane surface area, this also implies a reduction in surface area during the late stages of endocytosis, when *ψ*_max_ is large.

**Appendix 6—figure 1.**
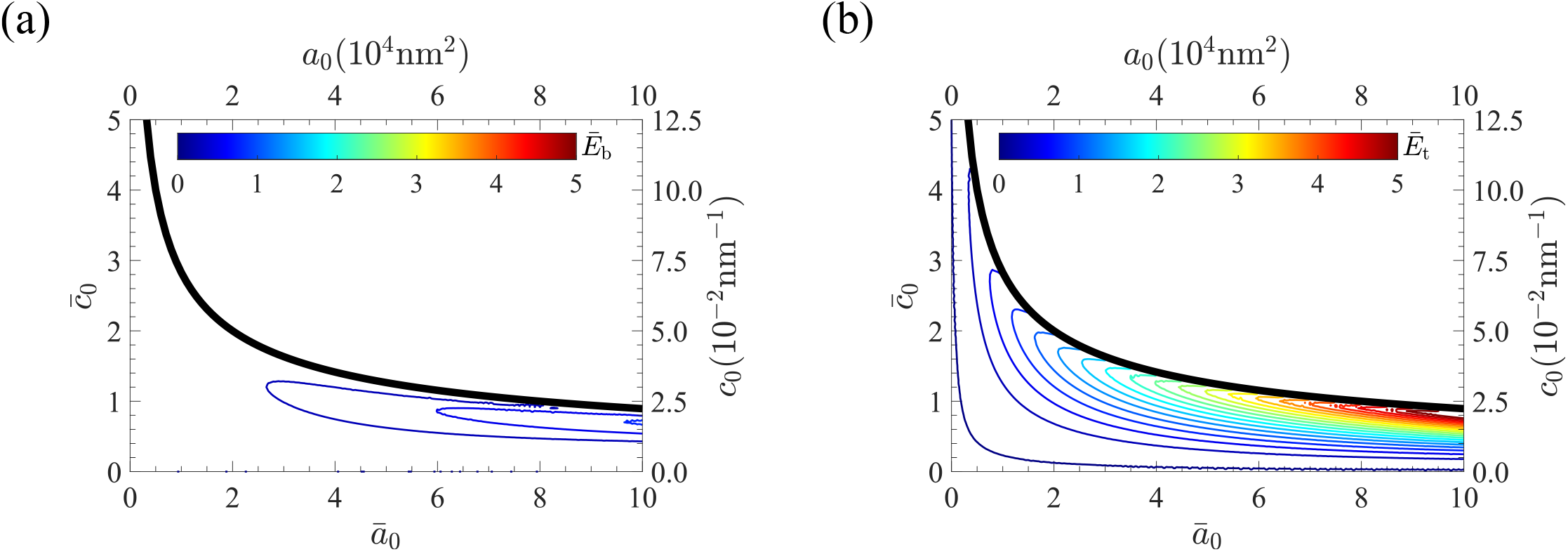
An appendix figure for ***Figure 4***. plots the energy landscape of membrane bending in (a) and the energy landscape of membrane tension in (b). A given 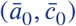 determines an unique value for both bending and tension energy, regardless of (*µ*, *ν*). The scales of colorbar in (a) and (b) are set the same to compare the contribution of the two energy terms. All colors, scales, ticks, etc., have the same meaning as in ***Figure 4***.

